# Pre-initiation and elongation structures of full-length La Crosse virus polymerase reveal functionally important conformational changes

**DOI:** 10.1101/2020.03.16.992792

**Authors:** Benoît Arragain, Grégory Effantin, Piotr Gerlach, Juan Reguera, Guy Schoehn, Stephen Cusack, Hélène Malet

**Affiliations:** Université Grenoble Alpes, CNRS, CEA, Institute for Structural Biology (IBS), F-38000, Grenoble, France; European Molecular Biology Laboratory, Grenoble, France; Department of Structural Cell Biology, Max Planck Institute of Biochemistry, Munich, Germany; Aix-Marseille Université, CNRS, INSERM, AFMB UMR 7257, 13288 Marseille, France

## Abstract

*Bunyavirales* is an order of segmented negative stranded RNA viruses comprising several life-threatening pathogens such as Lassa fever virus (*Arenaviridae*), Rift Valley Fever virus (*Phenuiviridae*) and La Crosse virus (LACV, *Peribunyaviridae*) against which neither specific treatment nor licenced vaccine is available. Replication and transcription of *Bunyavirales* genome constitute essential reactions of their viral cycle that are catalysed by the virally encoded RNA-dependent RNA polymerase or L protein. Here we describe the complete high-resolution cryo-EM structure of the full-length (FL) LACV-L protein. It reveals the presence of key C-terminal domains, notably the cap-binding domain that undergoes large movements related to its role in transcription initiation and a zinc-binding domain that displays a fold not previously observed. We capture the structure of LACV-L FL in two functionally relevant states, pre-initiation and elongation, that reveal large conformational changes inherent to its function. We uncover the coordinated movement of the polymerase priming loop, lid domain and C-terminal region required for the establishment of a ten-base-pair template-product RNA duplex before strand separation into respective exit tunnels. The revealed structural details and dynamics of functional elements will be instrumental for structure-based development of compounds that inhibit RNA synthesis by the polymerase.

## INTRODUCTION

Segmented negative stranded RNA viruses (sNSV) are divided into two major orders: *Bunyavirales*, that comprises more than 500 species classified in twelve families^1^, and *Articulavirales*, that include influenza virus from the *Orthomyxoviridae* family. Replication and transcription of sNSV viral genomic segments are performed by the virally encoded RNA-dependent RNA polymerase or L protein^2^. These processes are performed in the cytoplasm of infected cells for Bunyaviruses, whereas they occur in the nucleus for influenza virus^3, 4^. Replication generates full-length genome or antigenome copies (vRNA and cRNA respectively), whereas transcription produces capped viral mRNA that are recognized by the cellular translation machinery to produce viral proteins. Transcription is initiated by a “cap-snatching” mechanism, whereby host 5′ capped RNAs are bound by the L cap-binding domain (CBD), cleaved by the L endonuclease domain several nucleotides downstream, and then used to prime synthesis of mRNA^2, 5, 6^. Although the overall mechanism of transcription initiation is likely conserved between sNSVs, several elements suggest some divergences between viral families. First, the source of capped RNA differs. Whereas influenza polymerase interacts directly with the host polymerase II to snatch the caps of nascent transcripts in the nucleus^7^, it is currently unclear which and in what context cytoplasmic capped RNAs are accessed by bunyavirus polymerases and if the polymerase contains specific domains that would serve as platforms to interact with host capped-RNA-bound proteins. Second, the length of the host-derived capped RNA primer generated after cleavage by the endonuclease differs between families^5^, 0 to 7 in *Arenaviridae,* 10 to 18 in *Peribunyaviridae*, 10 to 14 in *Orthomyxoviridae,* suggesting differences in the relative position of the endonuclease, CBD and polymerase active site. Third, CBD localization within L proteins remains unclear for several viral families primarily due to the absence of a definitive motif for the cap-binding site and because of the high divergence in sequence between polymerases, particularly in their C-terminal region. Identification of the CBD in the C-terminal region of L proteins has however recently been achieved for both Rift Valley Fever virus (RVFV, *Phenuiviridae)* and California Academy of Science virus (CASV, *Arenaviridae*), thanks to the determination of isolated CBD domain structures^8, 9^.

To understand the detailed mechanisms of the replication and transcription reactions, structures of full-length polymerases are essential. Significant advances have recently been made on influenza polymerase with structures stalled at different steps of transcription now being available. They notably reveal that influenza CBD undergoes a 70° movement to bring the capped RNA from a position in which it can be cleaved by the endonuclease to a position in which it can enter into the polymerase active site^10–12^. A snapshot of transcription elongation has been captured revealing the presence of a nine-base-pair template-product RNA inside the active site cavity that is then separated into two single-stranded RNAs exiting through separated tunnels^12^. In comparison, structural information on *Bunyavirales* polymerase remains limited with only the structure of a C-terminally truncated construct of LACV polymerase (residues 1 to 1750, LACV-L_1-1750_) being currently available^13^. LACV-L_1-1750_ is composed of an N-terminal protruding endonuclease domain (residues 1-185) and a polymerase core comprising the active site (residues 186-1750). It was solved by X-ray crystallography in the pre-initiation state in complex with the 3′ and 5′ promoter ends. Both promoter ends bind in separate sequence-specific pockets away from the active site, respectively called “5′ end stem-loop pocket” and “3′ end pre-initiation pocket”. The LACV-L_1-1750_ structure also depicts the presence of an active site cavity with typical polymerase motifs as well as distinct template and product exit tunnels.

To reveal the structure of the C-terminal region of LACV-L and the overall architecture of the complete polymerase, we determined the structure of LACV-L FL by X-ray crystallography and high-resolution cryo-EM. We uncover the structure of LACV-L cap-binding domain, which contains specific insertions that relates to its interaction with the endonuclease. We find that the extreme C-terminal region is a zinc-binding domain that is absent in other structurally determined sNSV polymerases and may correspond to a host-protein interaction platform. We also capture snapshots of LACV polymerase in both pre-initiation and elongation-mimicking states, thereby revealing, amongst other conformational changes, the movement of the priming loop that unblocks the active site cavity and permits accommodation of the ten-base-pair template-product RNA characteristic of elongation.

## RESULTS

### Structure determination of LACV-L FL protein

LACV-L FL was expressed in insect cells, purified to homogeneity, and incubated with the 3′ and 5′ promoters following an adjusted version of the protocol described previously^13^ (**Supplementary Fig. 1a**, see methods). LACV-L FL was crystallized and its structure was solved at 4.0 Å resolution by molecular replacement using LACV-L_1-1750_ as a template, revealing two molecules in the asymmetric unit. There was clear extra density showing repositioning of the endonuclease and also for the previously missing C-terminal region, but the resolution was insufficient for building an accurate model (**Supplementary Fig. 1b, Supplementary Table 1)**. LACV-L FL was later on characterized by cryo-EM and resulted in a 3.0 Å resolution structure. A 2.28 million particle dataset was collected on a Titan Krios equipped with a K2 direct electron detector **(Supplement Fig. 2a)**. 2D and 3D classifications revealed that the C-terminal region of the polymerase is extremely flexible, and only a fraction of the dataset containing 0.37 million particles that displays a defined density for the C-terminal region was kept for further structural analysis **(Supplement Fig. 2b)**. The resulting “stable dataset” was further 3D classified resulting in the separation of two defined states: (i) the expected pre-initiation state and (ii) an elongation-mimicking state in which the complementary 3′ and 5′ vRNA formed a double stranded RNA that could be encapsulated within the active site cavity **(Fig. 1b and c, Supplement Fig. 2c)**. Even though only the “stable dataset” was used for 3D classification, the C-terminal region (residues 1752-2263) remained poorly defined due to flexibility. Advanced image analysis (see methods) was necessary in order to determine the structure of all C-terminal domains between 3.0 and 3.5 Å resolution (**Supplementary Fig. 2c, Supplementary Fig. 3, Supplementary Table 2)**. The complete model of LACV full-length polymerase was manually built and refined **(Fig. 1)**.

**Figure 1:**
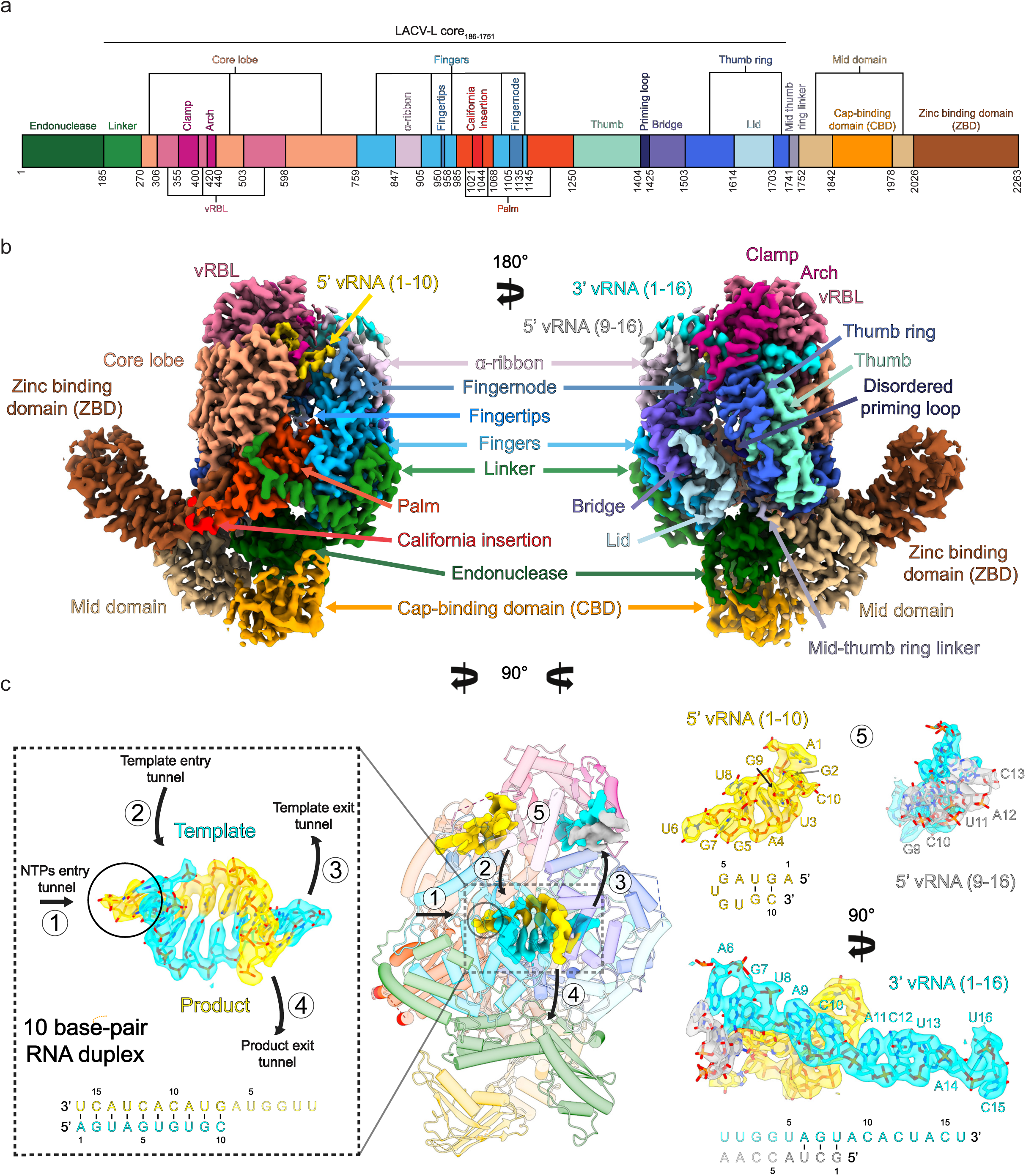
Cryo-EM structures of LACV-L FL. **a**, Schematic representation of LACV-L FL domain structure. **b**, Two views of LACV-L FL cryo-EM map at pre-initiation stage. A composite cryo-EM map was assembled from individual maps of the (i) LACV-L core, endonuclease and mid domain, (ii) mid domain and CBD and (iii) mid domain and ZBD. Domains and RNAs are indicated with arrows and colored as in **a**. **c**, Cartoon representation of semi-transparent LACV-L FL cryo-EM structure at elongation-mimicking stage rotated of 90° compared to **b**. Close-up views of the Coulomb-potentials and models of all RNAs visible in the elongation-mimicking map. The position of the four tunnels are shown and numbered from 1 to 4. The 5′ end vRNA (1-10) in its “5’ end stem-loop loop pocket” is displayed in yellow, the 3’ vRNA (1-16) in its “3′ end pre-initiation pocket” is shown in cyan, the 5’ vRNA (9-16) that hybridizes with the 3’ vRNA (1-16) is colored in light grey. The RNA that mimics the template and product are shown in cyan and yellow respectively. The sequence and secondary structures of nucleic acid moieties in each complex is displayed.

### Overall structure of LACV-L FL

The X-ray and cryo-EM structures reveal the same overall arrangement of LACV-L FL. The polymerase core (residues 186-1751) is conserved compared to the LACV-L_1-1750_ construct (RMSD of 0.474 Å on 1187 Cα) but the endonuclease domain undergoes a large rotational movement of 180° (**Supplementary Fig. 4**). LACV-L FL C-terminal region (1752-2263) protrudes away from the core and forms an elongated arc-shaped structure that is supported and stabilized by a β-hairpin strut (residues 2084-2102) that bridges to the core (**Fig. 1b, Fig. 2**). At one end of the C-terminal domain is the CBD (residues 1842-1977) and at the other end the zinc-binding domain (ZBD, residues 2026-2263) (**Fig. 1b, Fig. 2a**). They are both connected to the mid-domain (residues 1752-1841 and 1978-2025) (**Fig. 1b, Fig. 2a**).

**Figure 2:**
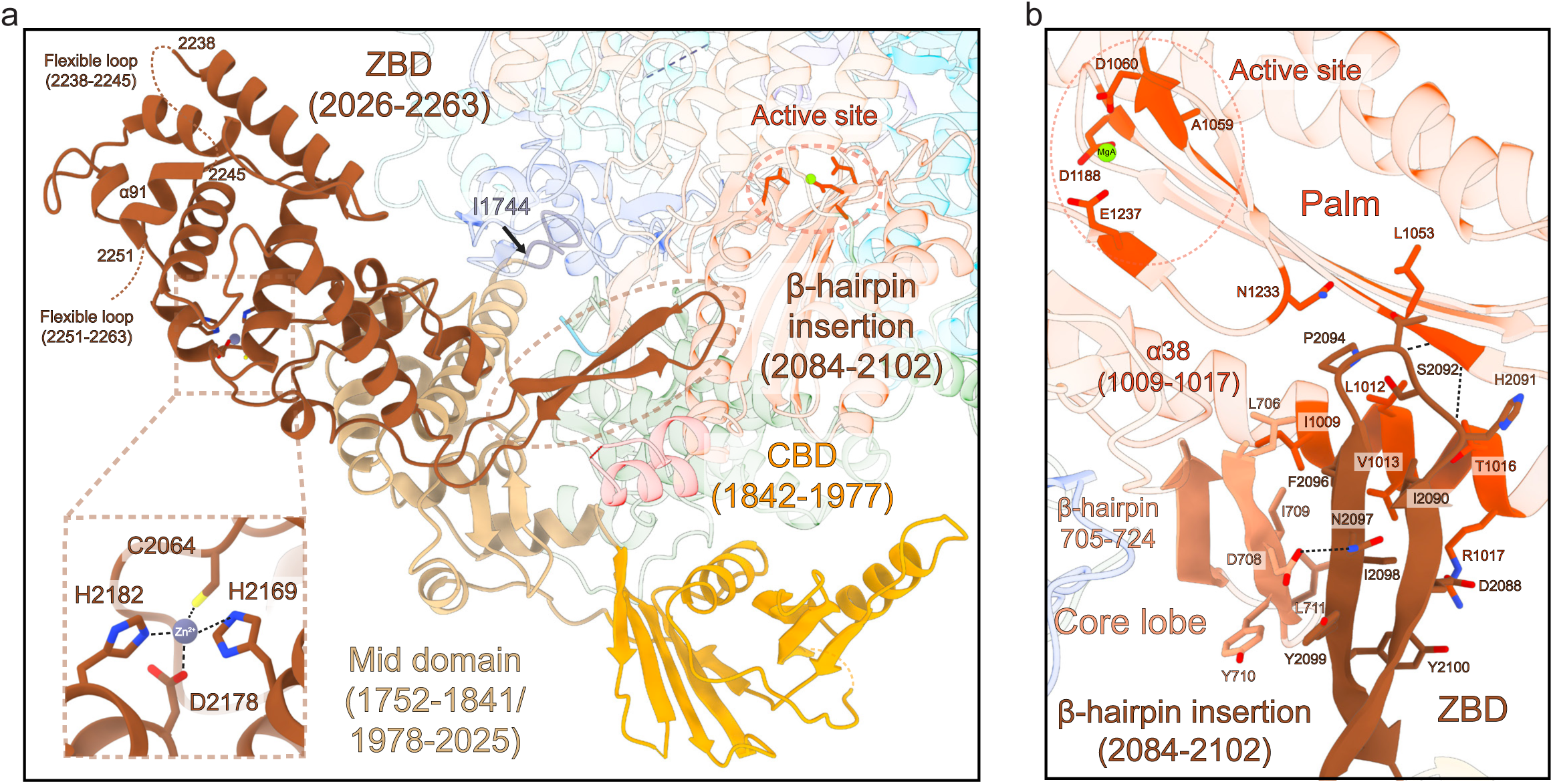
C-terminal region of LACV-L FL. **a**, Structure of LACV-L C-terminal region. The ZBD β-hairpin that protrudes towards the core is surrounded by a dotted line. Position of the last α-helix 91 is indicated. Close-up view on the coordination of the presumed zinc ion from the ZBD. **b**, Close-up view of the ZBD β-hairpin. Residues from the core lobe β-hairpin and the palm domain that interacts with the ZBD β-hairpin are indicated. The active site is surrounded by a dotted line.

### Structure and mobility of the cap-binding domain of LACV-L

The CBD is composed of a five-stranded anti-parallel β-sheet (β34, β35, β36, β37, β41) packed against the α-helix 77 that is flanked by a three-stranded antiparallel β-sheet (β38, β39, β40), the α-helix 78 and long loops (**Fig. 3a)**. There is a disordered loop between the first two strands of the CBD five-stranded β-sheet (**Fig. 3 a,b**) that contains a number of residues highly conserved in all *Peribunyaviridae* L proteins, although there is no density for them. The m^7^GTP binding sites of RVFV and influenza virus CBD^8, 14^ which share the same overall fold are located in an equivalent loop (**Supplementary Fig. 5**). The m^7^GTP cap-binding site of LACV L can therefore be predicted to be composed of W1847 and W1850 that would sandwich the m^7^GTP, supported by Q1851 and R1854 that would respectively interact with guanine moiety and phosphates (**Fig. 3b, Supplementary Fig. 5**). This suggests a conserved mode of m^7^GTP interaction mediated by functionally equivalent residues without any overall sequence conservation.

**Figure 3:**
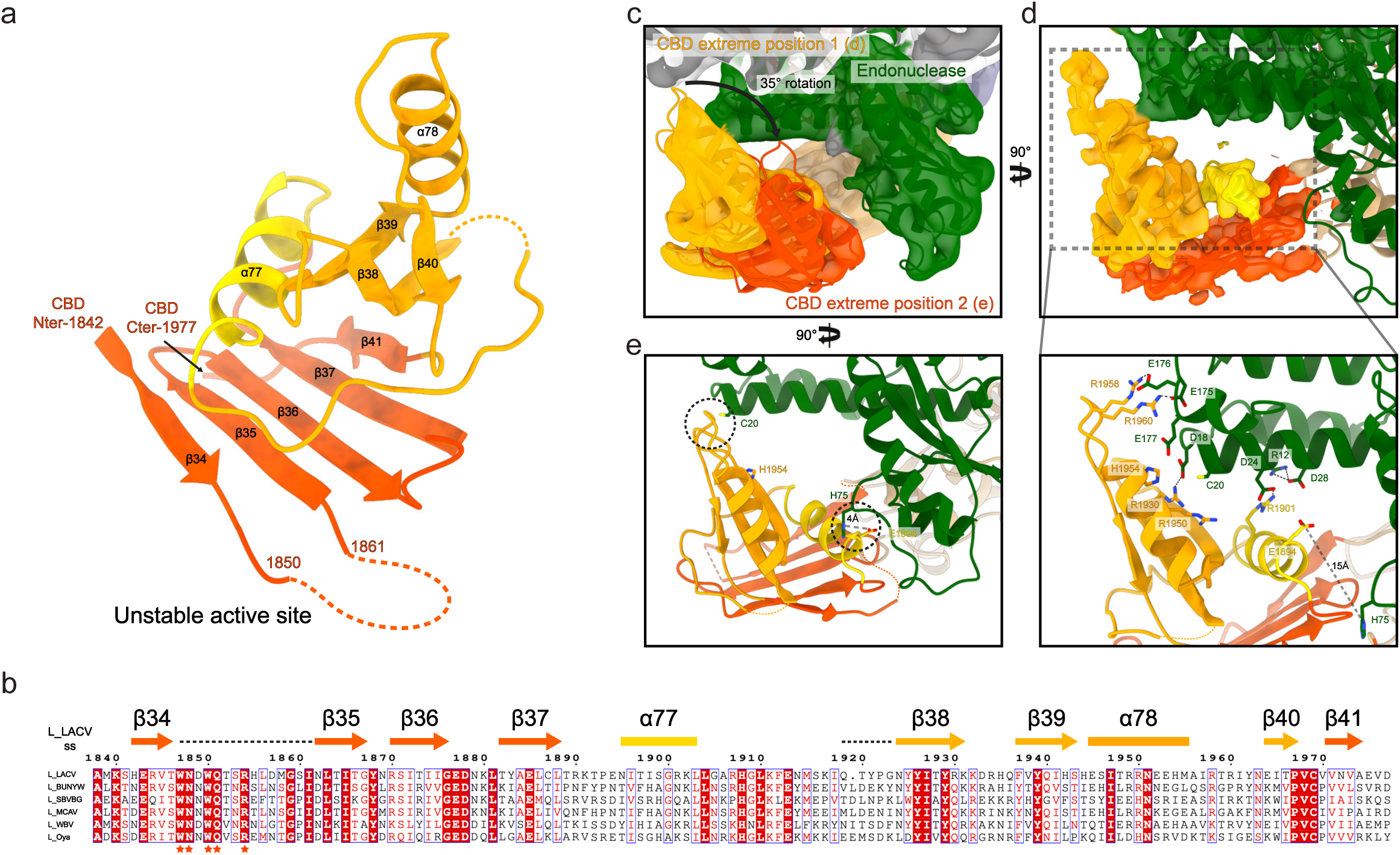
LACV-L CBD structure and its interaction with the endonuclease. **a**, LACV-L CBD atomic model. Secondary structures are shown. The fold conserved with other sNSV CBD is shown in red and yellow. LACV CBD insertion is shown in orange. The missing loop comprising the active site is shown as a dotted line. **b**, Multiple alignment of six *Peribunyaviridae* CBD: La Crosse virus (LACV), Bunyamwera virus (BUNYW), Schmallenberg (SBVBG), Macaua virus (MCAV), Wolkberg virus (WBV) and Oya virus. Secondary structures of the CBD LACV-L are shown and colored as in **a**. Missing residues of LACV-L CBD are presented as dotted lines. Conserved residues of *Peribunyaviridae* CBD active site motif (WNxWQxxR) are shown as orange stars. **c**, Cryo-EM 3D classes corresponding to LACV-L CBD extreme position 1 and 2 are superimposed. Their CBD are respectively shown in gold and red. The endonuclease domain is shown in green. LACV-L core is shown in grey. **d**, Overview and close-up view of the CBD/endonuclease domain interactions in the extreme position 1 are displayed. Interacting residues are identified and labelled. CBD coloring is the same as in **a**. **e**, Overview of the CBD/endo-nuclease domain interaction in the extreme position 2 is shown. Interacting residues are identified and labelled. CBD side chain positions remain however hypothetical due to the CBD EM map resolution in extreme position 2.

The CBD is rotationally mobile as visualized in a 3D variability analysis of the dataset (**Supplementary Video 1**). Its large movements are enabled by the conformationally stable mid domain that acts as a central hub mediating contacts between the core, the CBD and the ZBD (**Fig 2a**). Several CBD positions can be separated by 3D classification (**Supplementary Fig. 2c**, see methods) and a rotation of 35° is visible between the two extreme positions (**Fig. 3c**). In the extreme position 1, residues 12-28 and 175-178 of the endonuclease domain interact with residues E1894, R1901, R1930, 1950-1960 of the CBD mainly through electrostatic interactions (**Fig. 2d**). In the extreme position 2, the contacts between the CBD and the rest of the polymerase are rather sparse, explaining its instability (**Fig. 2e**). The only interactions are mediated by the loop 1932-1936 of the CBD that is proximal to C20 of the endonuclease domain, and the residue E1894 of the CBD that is close to the H75 of the endonuclease domain (**Fig. 2e**).

### The zinc-binding domain: a protruding region that connects to the core through a β-hairpin strut

The C-terminal extremity of LACV-L is a α-helical domain with a long protruding β- hairpin (**Fig. 2a,b**). Its two sub-domains of equivalent size surround a metal ion that is coordinated by four residues highly conserved amongst peribunyaviruses: C2064, H2169, D2178 and H2182, suggesting that it is a zinc ion (**Fig. 2a, Supplementary Fig. 10**). The overall topology of the ZBD has not been previously observed according to a DALI search^15^. This domain protrudes out of the polymerase, suggesting that it could be extremely mobile. This appears to be the case for many of the particles, impeding their use in structure determination of this domain (**Supplementary Fig. 2c**). However, in the particles used for high-resolution determination of the C-terminal region, the above-mentioned long protruding β-hairpin (residues 2084-2102, **Fig. 2a**) stabilizes the ZBD with the core via the formation of a four-stranded antiparallel β-sheet composed of the ZBD β-hairpin and a β-hairpin from the core lobe (residues 705-724) (**Fig. 2b**). Interestingly, that region was not visible in the LACV-L_1-1750_ electron density and is only structured in the presence of the ZBD β-hairpin. In addition, the ZBD β-hairpin makes several hydrophobic interactions with residues 1009-1017, L1053 and N1233 of the palm domain, proximal to the polymerase active site (**Fig. 2b**). The extreme C-terminal α-helix 91 of the ZBD (**Fig. 2a and Supplementary Fig. 1c**) is connected to the rest of the domain via a long flexible loop permitting large movements. In the crystal structure, the α-helix 91 protrudes away to bind to a hydrophobic pocket present in the ZBD of the second polymerase of the asymmetric unit, forming a domain-swap dimer (**Supplementary Fig. 1c**). In the cryo-EM map, the polymerase is monomeric and the α-helix 91 folds back into the same hydrophobic pocket of the ZBD (**Fig. 2a)**.

### Elongation-mimicking state

Based on the RNA promoter sequences the polymerase was incubated with, we were expecting to obtain only the structure at pre-initiation state. However, extensive 3D classification resulted in identification of an alternative RNA-bound subset of particles in which a remarkable ten-base-pair duplex is visible in the active site cavity (**Supplementary Fig. 2c**). The structure, mimicking an elongation state with a bound product-template duplex, is determined at 3.0 Å resolution, enabling to distinguish unambiguously purine and pyrimidine bases (**Fig. 4a**). It can thus be deduced that the RNA duplex corresponds to the hybridisation between the nearly complementary 5′ and 3′ promoter ends (5′p-(1)AGUAGUGUGC(10) and 3′OH-(16)UCAUCACAUG(7)), corresponding to nucleotides 1-10 for the 5′ and 7-16 for the 3′ (**Fig. 4a,c,d**). Visualisation of this state shows that a small fraction of the “stable dataset subset” (59,590 particles out of 370,497, **Supplementary Fig. 2c**) was able, in the *in vitro* conditions used and with 3′ and 5′ promoters in excess, to internalize the promoter duplex in the active site cavity. This is in addition to the 3′ and 5′ RNA promoters being also bound in their “3′ end pre-initiation pocket” and “5′ stem-loop pocket” respectively, in positions identical to the ones observed at pre-initiation, showing the RNA-binding compatibility between all these separate RNA-binding sites (**Fig. 1c**). Although not a true elongation state, the structure obtained fortuitously mimics this state and gives insight into the mechanisms of (i) RNA binding in the active site cavity and (ii) template-product separation after formation of a ten-base-pair double-stranded RNA in the active site cavity.

**Figure 4:**
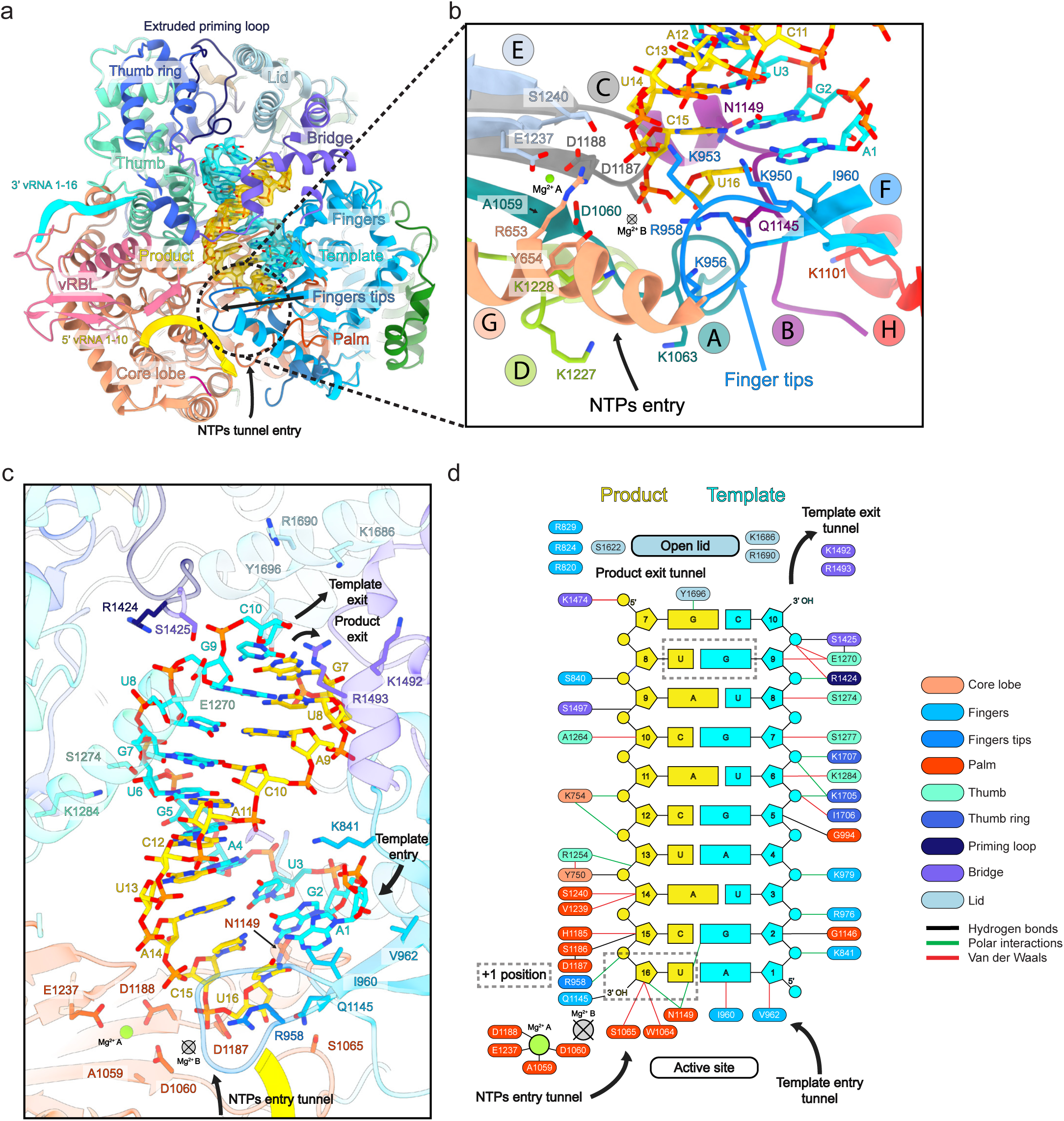
Cryo-EM structure of the LACV-L FL at elongation-mimicking stage. **a**, Cut-away view of the LACV-L FL at elongation-mimicking stage. Its orientation corresponds to a 90° rotation compared to Fig.1 b left (top view visualization of Fig.1 b left). The main domains are depicted and colored as in Figure 1. The Coulomb potential map of the ten-base-pair product-template RNA is shown in gold for the product (3’OH-UCAUCACAUG, nucleotides 7 to 16) and cyan for the template (5’p-AGUAGUGUGC, nucleotides 1 to 10). The extruded priming loop is shown in dark blue. The active site position is indicated as a dotted line. **b**, View of the LACV-L FL active site showing the conserved RNA-dependent RNA polymerase (RdRp) functional motifs A to H (G and H are only conserved in sNSV polymerases). They are respectively colored turquoise, purple, grey, light green, light blue, blue, beige and red for A to H. Template- and product-mimicking RNA are colored as in **a**. Metal A, presumed to correspond to a magnesium ion, is shown as a green sphere. Metal B is absent but its supposed position obtained by superimposition with active RdRps is shown as a crossed circle. **c**, Interactions of the ten-base-pair product-template RNA in LACV-L active site cavity. Principal residues from the active site in palm domain (A1059, D1060, S1065, D1187, D1188, E1237), fingertips (R958), fingers (K841, I960, V962, Q1145), bridge (R1424, S1425, K1492, R1493), thumb (S1274, K1284) and lid (K1686, R1690, Y1696) are displayed. NTPs entry, template entry, template entry/exit tunnel directions are shown. Position of ions is shown as in **b**. Nucleotides are labelled according to RNA promoter sequence. **d**, Schematic representation of RNA-protein contacts in the active site cavity. Residues are colored according the domain to which they belong, the same domain coloring as in **a** is used. Template and product RNA are numbered according to their position in the 5′ end promoter (5′p-AGUAGUGUGC, nucleotides 1 to 10) and the 3′ end promoter (3’OH-UCAUCACAUG, nucleotides 7 to 16). The U-G mismatch that is due to the non-perfect complementarity between the 5′ and 3′ promoters is surrounded by a dotted rectangle. Interaction type are color coded as indicated. Ions are shown as in **b** and **c**. Active site and lid domain positions are indicated. Nucleotide U16 corresponds to the nucleotide in position +1 of the product and is identified as such.

The template-product duplex interacts in the active site chamber primarily through its backbone as well as bases with a large number of residues of the polymerase making both van der Waals and polar interactions (**Fig. 4c,d**). The template-mimicking RNA that is proximal to the active site (nucleotides 1, 2 and 3) interacts with the finger domain, the central part (nucleotides 4, 5) binds to the palm, while the distal template-mimicking RNA (nucleotides 6-10) interacts with the thumb and the thumb ring domains (**Fig. 4a,c,d**). The proximal part of the product-mimicking RNA (nucleotides 14-16) is surrounded by the palm domain, the central part of the product (nucleotides 10-13) interacts with the core and the core lobe, while the distal part of the product-mimicking RNA (nucleotides 7-9) mainly binds to the bridge and the finger domains (**Fig. 4a,c,d**). The LACV-L catalytic core shares with other viral RNA-dependant RNA-polymerases the six conserved structural motifs (A-F)^16^ (**Fig. 4b**). In addition, motifs G and H that are specific to sNSV polymerases are also visible^13^ (**Fig. 4b**). The polymerase conformation mimics an elongation post-incorporation, pre-translocation step in which an incoming nucleotide would just have been incorporated into the product. The active site is slightly more open than what is normally observed after incorporation, probably due to the absence of pyrophosphate to be released. In order to be active a polymerase normally binds to two divalent positively charged ions (metal A and B) that control nucleotide addition^16^. In the LACV-L elongation-mimicking structure, there is a presumed Mg^2+^ ion coordinated by residues D1188 (motif C), E1237 (motif E) and the carbonyl oxygen of A1059 (motif A) (**Fig. 4b**). It corresponds to the typical position for metal A in the inactive open state. The Mg^2+^ ion that would corresponds to metal B is absent. Based on architectural similarities of viral RNA polymerases^16^, local reconfigurations of motifs A and C will be required to re-orientate the active site triad D1060 (motif A), D1187 (motif C) and D1188 (motif C) to correctly bind the two metals in a catalytic configuration (**Fig. 4b**). The other motifs are already in an active conformation. Fingertips residues R958 and I960 respectively stack on the bases of the product and template nucleotides in the +1 position, thereby stabilizing them (**Fig. 4b**). The motif B loop, which is implicated in the selection of the correct nucleotide to be incorporated, adopts a conformation compatible with the post-incorporation state. Its residue Q1145 contacts the +1 position nucleotide base of the last incorporated product nucleotide, while residue N1149 interacts with the 2′ hydroxyl group of the template nucleotide (**Fig. 4b**). In summary, the elongation-mimicking structure would just need local closure coupled to local reconfiguration of motifs A and C to represent the post-incorporation pre-translocation elongation step containing a ten-base-pair template-product RNA in the active site chamber.

### Conformational changes between pre-initiation and elongation

Comparison between the pre-initiation and elongation-mimicking states reveals key movements of the L protein in action. The priming loop (residues 1404-1424) is an essential element that stabilises the first nucleotide to be incorporated in the product during replication initiation stage. In the pre-initiation structure, it protrudes towards the active site but is disordered (**Fig. 5a)**. As part of the initiation to elongation transition, it extrudes from the active site via the template exit tunnel, thereby freeing space for the ten-base-pair RNA to fit in the active site chamber (**Fig. 5b)**. The fully ordered and extruded priming loop is located on the surface of the thumb ring and lid domains (**Fig. 5b**) with which it interacts mainly through hydrophobic contacts involving residues V1572, Y1576, A1751, M1753, N1658, L1661 (**Fig. 5d**). Interestingly, the priming loop movement is coupled with the reorganisation of mid-domain residues 1752 to 1761 from an α-helix to an extended loop. This results in a modification of the mid-thumb-ring linker: it encompasses residues 1741 to 1751 at pre-initiation and extends to residues 1741 to 1761 at elongation. As a result, the mid-thumb-ring linker loop extremity, comprising residues 1750-1753 at elongation, is displaced by 8 Å between pre-initiation and elongation (**Fig. 5d**) and interacts with the priming loop residues 1414-1418 mainly through hydrophobic contacts (**Fig. 5d**).

**Figure 5:**
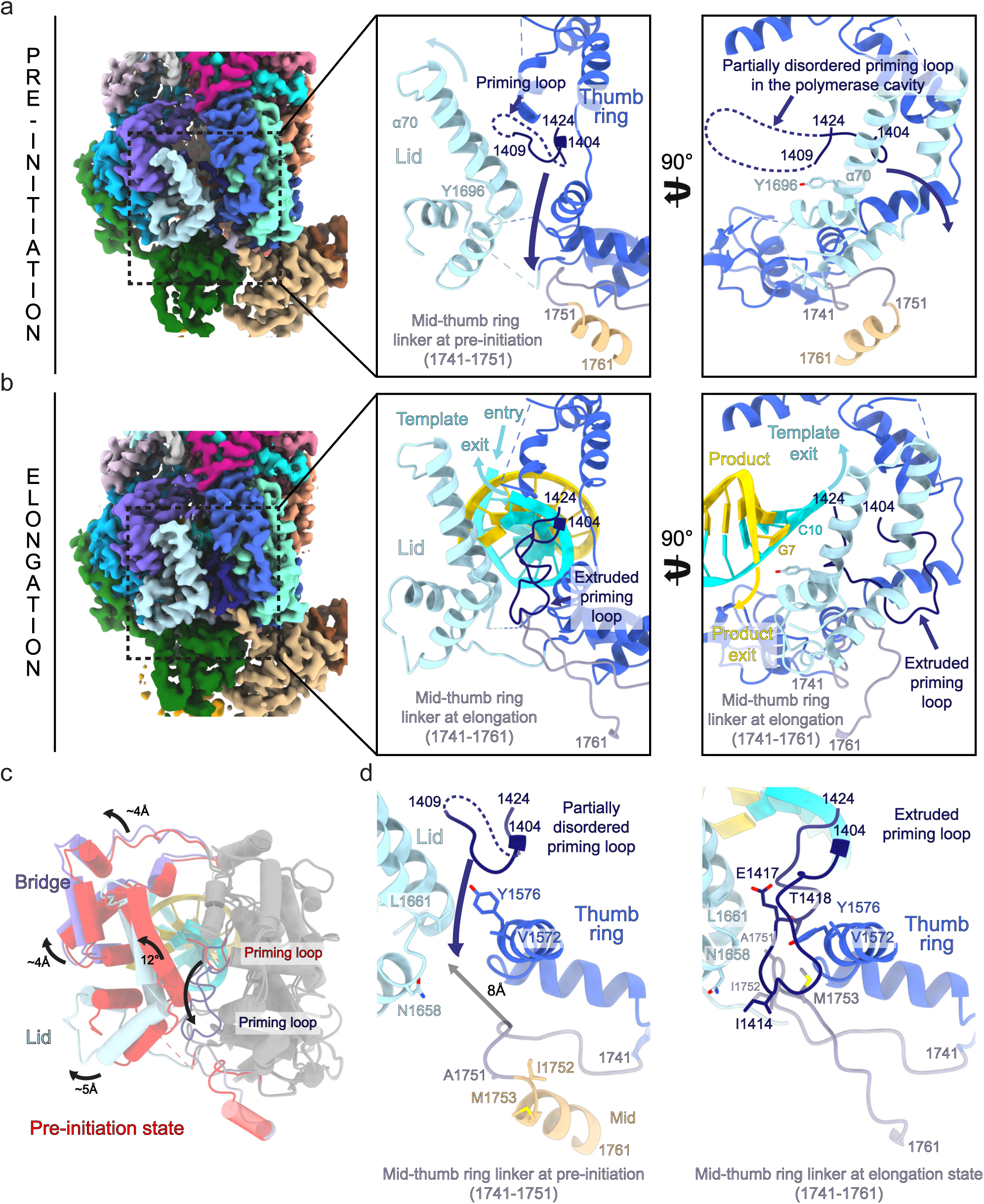
Conformational changes between pre-initiation and elongation-mimicking states. **a**, LACV-L FL in pre-initiation state. Left panel: cryo-EM map colored as in Figure 1. Middle panel: close-up view. Right panel: close-up view rotated 90° compared to the middle panel. For more clarity, only the lid, thumb ring and mid domains are shown in the middle and right panels and are colored in light blue, blue and beige. The mid-thumb ring linker (1741-1751) is shown in purple. The priming loop is shown in dark blue. All these elements are labelled. The lid domain is closed and the residue Y1696 from the α-helix 70 points towards the cavity. The priming loop is disordered between residues 1410 and 1423 (dotted line). Priming loop movement occurring during elongation is depicted with a dark blue arrow. **b**, LACV-L FL in elongation-mimicking state. Equivalent panels as in a are shown with domains and loops colored as in **a**. The mid-thumb ring linker contains residues 1741 to 1761 at elongation. The template tunnel entry and exit, the product exit are shown. The lid domain is open and Y1696 from the α-helix 70 interacts with the product-mimicking RNA. The priming loop is ordered and interacts with the lid, the thumb ring and the mid-thumb ring linker. **c**, Superimposition of the thumb and thumb-ring (shown in grey) in pre-initiation and elongation-mimicking state of LACV-L FL. The bridge and lid in pre-initiation are shown in red. The bridge and lid in elongation-mimicking state are shown in purple and light blue. Their rotation between the two states is labelled. Movement of the priming loop between the two states is shown with an arrow. **d**, Priming loop and mid-thumb ring linker at pre-initiation (left) and elongation (right). Numbering of both elements are indicated. Residues from the lid and thumb-ring that interact with the priming loop at pre-initiation and/or elongation are shown. The 8 Å displacement between the mid-thumb ring linker extremity at pre-initiation and elongation is indicated with an arrow.

The transition from pre-initiation to elongation is also coupled to coordinated domain movements. The lid domain moves by 12° compared to the thumb ring, resulting in the opening of the template and the product exit tunnels (**Fig. 5c)**. Separation of the template-product RNA duplex is made possible by the α-helix 70 of the lid domain that faces the distal part of the double-stranded RNA, and in particular its residue Y1696 that interacts with the 5′ end product nucleotide, thereby forcing strand-separation of the RNA duplex (**Fig. 4b)**. Domain movements occurring between pre-initiation and elongation are nicely captured by a 3D variability analysis of the dataset (**Supplementary Video 2**). It reveals a coordinated rotation of the endonuclease and the C-terminal region compared to the core, using the mid domain as a hinge, and resulting in 4.5 Å and 8 Å displacement of the endonuclease and the ZBD respectively (**Supplementary Video 2**).

## DISCUSSION

### Overall comparison between C-terminal domains of negative stranded RNA virus polymerases

The structure presented here reveals the organization of the entire LACV-L protein. The newly described C-terminal domains can be compared with equivalent parts of *Phenuiviridae*, *Arenaviridae* and *Orthomyxoviridae* polymerases.

LACV-L CBD shares a conserved minimal fold with equivalent structures from RVFV (*Phenuiviridae*)^8^, CASV (*Arenaviridae*)^9^ and influenza virus (*Orthomyxoviridae*)^14^, consisting in an antiparallel β-sheet stacked against an α-helix (**Supplementary Figure 5**). In addition, LACV-L CBD contains an insertion consisting of the three-stranded β-sheet (β38, β39, β40), the α-helix 78 and charged loops (1932-1936 and 1956-1963). This insertion is likely to be related to the CBD role in LACV-L transcription initiation as it is implicated in interactions with the endonuclease (**Fig. 3d**).

LACV C-terminal region also contains a mid-domain and a ZBD. The mid-domain fold is conserved between *Orthomyxoviridae* and *Arenaviridae* polymerases^9, 11^ (**Supplementary Figure 6b**), whereas the distal C-terminal region of the polymerase differs between the three families. *Orthomyxoviridae* and *Arenaviridae* respectively have a PB2 627-domain and D1-III domain that are structurally related^9, 11^, whereas the ZBD present in LACV has a different fold. This suggests that *Orthomyxoviridae* and *Arenaviridae* polymerases are more closely related (**Supplementary Figure 6c**). The presence of the ZBD or 627-domain in the C-terminal region in a protruding position compared to the polymerase core suggests that, as for the 627-domain^17^, LACV ZBD may play a role in host protein interaction. This leads us to hypothesize that it might act as a platform to recruit cytoplasmic proteins recognising capped RNA or be involved in replication-related activities. It might as well interact with host translation factors, thus mediating the transcription-translation coupling observed in *Bunyavirales*^5, 18^.

### Comparison of CBD movements occurring in LACV and influenza virus polymerases

Particles of the LACV-L dataset have variable CBD positions, with extreme conformations being 35° apart (**Fig 3.c, Supplementary Fig. 7a,b, Supplementary Video 1)**. The observed movements are likely to be correlated to the CBD function at the pre-initiation stage. In its extreme position 1, the CBD active site is exposed towards the exterior, in a favourable position to interact with the cellular capped RNA (**Supplementary Fig. 7a, Supplementary Video 1**). Position 2 would bring the capped RNA closer to the endonuclease active site (**Supplementary Fig. 7b**). Interestingly, the position of both the endonuclease and the CBD compared to the core differ from the arrangement observed in influenza virus polymerase (**Supplementary Fig. 7c,d**). LACV-L endonuclease domain is very close to the capped RNA entry tunnel (that also corresponds to the product exit tunnel), whereas in influenza virus polymerase the endonuclease is located away from the equivalent tunnel. In consequence, a 70° movement of influenza CBD brings the capped RNA from a position in which it can be cleaved by the endonuclease to a position in which capped RNA can enter the active site (**Supplementary Fig. 7c,d**). Even if the 35° movement of LACV-L CBD between its extreme positions 1 and 2 brings the RNA closer to the endonuclease active site, additional movements would then be necessary to bring the capped mRNA primer in the active site. This suggests some divergence in the cap-snatching mechanism of the two viruses. One can also speculate that the endonuclease domain would need to relocate after transcription initiation and move away from the newly transcribed RNA in order to prevent transcript degradation. The insight provided here constitutes a basis to further address the exact mechanisms underlying LACV-L transcription initiation in greater detail in the future.

### Comparison between LACV and influenza polymerase elongation states and conformational changes between pre-initiation and elongation

LACV-L FL cryo-EM structures reveal significant movements between pre-initiation and elongation stages. Certain aspects are reminiscent to those reported for influenza virus polymerase^12^. Their active site cavities with bound template-product duplex are remarkably similar. The presence of 9-bp dsRNA in influenza virus polymerase elongation state^12^ versus 10-bp dsRNA in LACV-L comes from the fact they are in slightly different stages, influenza virus polymerase being in pre-incorporation state, meaning the last nucleotide has not yet been incorporated, whereas LACV-L structure mimics a post-incorporation elongation state and therefore contains an additional base-pair (**Supplementary Fig.8**). The mechanism of template-product separation by the lid domain is also remarkably conserved with equivalent helix of the lid implicated^12^. The tyrosine that prevents double-strand continuation strand differs however slightly in position: Y1696 of LACV-L interacts with the RNA product whereas Y207 of influenza PB2 stacks with the last nucleotide of the template (**Supplementary Fig.8**). Movements occurring between pre-initiation and elongation also differ between the two viral polymerases. For instance, their priming loop takes a completely different position and organization when it extrudes from the active site (**Supplementary Fig.9**). Influenza virus polymerase priming loop, which is 35-amino acid long, has 17 residues extruded into the solvent in a disordered loop, and projects towards the PB1-PB2 interface helical bundle^12^ (**Supplementary Fig.9b**). LACV-L priming loop, which is 20-amino acid long, orders itself at the surface of thumb ring and lid domains (**Supplementary Fig.9a**). Its extrusion is coupled with a *Peribunyaviridae* specific rearrangement of the α-helix 72 of the mid domain into a loop that extends the mid/thumb ring linker, enabling an interaction between the priming loop and the mid/thumb ring linker (**Fig. 5d**). Conformational changes of the whole C-terminal region upon the transition from pre-initiation to elongation visualized in the 3D variability analysis have also not been observed in influenza polymerase (**Supplementary Video 2**).

Altogether, the complete structure of LACV-L FL captured in pre-initiation and elongation-mimicking states and the rotational mobility of the CBD provide mechanistic insight into bunyaviral transcription. This, reinforced by the atomic details of the polymerase active site, establish a firm basis for future structure-based drug design that could target essential activities or critical conformation changes of *Peribunyaviridae*-L proteins.

## METHODS

### Cloning, Expression and Purification

Sequence-optimized synthetic DNA encoding a N-terminal his-tag, a TEV protease recognition site and the LACV-L (strain LACV/mosquito/1978, GenBank: EF485038.1, UniProt: A5HC98) was synthetized (Geneart) and cloned into a pFastBac1 vector between NdeI and NotI restriction sites. The LACV-L-expressing baculovirus was generated via the standard Bac-to-Bac method (Invitrogen). For large scale expression, Hi5 cells at 0.5×10^6^ cells/mL concentration were infected by adding 0.1% of virus. Expression was stopped 72h after the day of proliferation arrest. The cells were disrupted by sonication for 3 min (10 sec ON, 20 sec OFF, 50% amplitude) on ice in lysis buffer (50 mM Tris-HCl pH 8, 500 mM NaCl, 20 mM Imidazole, 0.5 mM TCEP, 10% glycerol) with EDTA free protease inhibitor complex. After lysate centrifugation at 20,000 rpm during 45 min at 4°C, protein from the soluble faction was precipitated using (NH_4_)_2_SO_4_ at 0.5 mg/ml and centrifuged at 30,000 rpm for 45 min, 4°C. Supernatant was discarded, proteins were resuspended back in the same volume of lysis buffer and centrifuged at 20,000 rpm during 45 min at 4°C. LACV-L was purified from the supernatant by nickel ion affinity chromatography after a wash step using 50 mM Tris-HCl pH 8, 1M NaCl, 20 mM Imidazole, 0.5 mM TCEP, 10% glycerol and eluted using initial lysis buffer supplemented by 300 mM Imidazole. LACV-L fractions were pooled and dialyzed 1h at 4°C in heparin loading buffer (50 mM Tris-HCl pH 8, 250 mM NaCl, 0.5 mM TCEP, 10% glycerol). Proteins were loaded on heparin column and eluted using 50 mM Tris-HCl pH 8, 1 M NaCl, 0.5 mM TCEP, 5% glycerol. LACV-L was then mixed in a 1:3 molar ratio with both 3′ (1-16) and 5′ (9-16) vRNA ends oligonucleotides - which had been pre-annealed by heating at 95°C for 2-5 min followed by cooling down on bench at RT temperature. During overnight dialysis at 4°C in a gel filtration buffer (20 mM Tris-HCl pH 8, 150 mM NaCl, 2 mM TCEP) LACV-L formed a complex with vRNA, which was ultimately resolved on the S200 size exclusion chromatography column.

Before freezing and storing at −80°C, LACV-L bound to 3′ (1-16) and 5′ (9-16) vRNA were mixed with 5′ (1-10) vRNA hook in a 1:2 molar ratio.

### Crystallisation and X-ray crystallography

For crystallisation, LACV-L in complex with pre-annealed 3′ (1-16) and 5′ (9-16) vRNA was concentrated to 5 mg/ml. The 5′ (1-10) vRNA end was later soaked into crystals. Initial hits were dense and round precipitates that appeared in 100 mM Tris pH 8.0, 100 mM NaCl, and 8% PEG 4000. Upon manual reproduction in hanging drops, they grew as thin hexagonal plates, but were soft and fragile and diffracted only to ∼8 Å. To improve the resolution, crystals were soaked in a stepwise manner with increasing concentration of the glycerol cryo-protectant, reaching 30%. Diffraction data were collected at the European Synchrotron Radiation Facility (ESRF), using a helical collection strategy and maximum transmission of the ID29 beamline. Crystals are of space-group *C*2, diffracting at best to a maximum resolution of 4.0 Å. Data were integrated with STARANISO^19^ to account for the anisotropy (**Supplementary Table 1**). The structure was solved with PHASER^20^ using LACV-L_1-1750_ (PDB code: 5AMQ)^13^ as a model after removal of the endonuclease. There are two L protein complexes per asymmetric unit (**Supplementary Fig.1b**). The initial map after molecular replacement revealed that the core of the L protein and bound RNA were little changed but there was clear density for the endonuclease in a new position. In addition, there was extra density for the C-previously missing C-terminal domain. This density was improved by multi-crystal and non-crystallographic 2-fold averaging using PHENIX^21^. Based on secondary structures predicted from an extensive multiple sequence alignment of *Peribunyaviridae* L proteins^22^, it was possible to build an approximate model of much of the C-terminal domain (which is largely helical), except the cap-binding domain for which there is no density, including identification of the zinc-binding site co-ordinated by highly conserved cysteine and histidine residues. The two C-terminal domains from the two complexes in the asymmetric unit interact around a non-crystallographic 2-fold axis in such a way that the extreme terminal helix of one packs against the C-terminal domain of the other, forming a domain swapped dimer. When the accurate structure of the C-terminal domain was obtained by cryo-EM, the X-ray model could be improved (**Supplementary Fig.1c, Supplementary Table 1**).

### Electron microscopy

For cryo-EM grid preparation, UltraAuFoil grids 300 mesh, R 1.2/1.3 were negatively glow-discharged at 30 mA for 1 min. 3.5 µl of the sample were applied on the grids and excess solution was blotted away with a Vitrobot Mark IV (FEI) (blot time: 2 sec, blot force: 1, 100% humidity, 20°C), before plunge-freezing in liquid ethane. Grid screening and cryo-EM initial datasets were collected on a 200kV Thermofischer Glacios microscope equipped with a Falcon II direct electron detector.

A high quality cryo-EM grid pre-screened on a 200kV Thermofischer Glacios microscope was used to collect data on a Thermofischer Titan Krios G3 operated at 300 kV equipped with a Gatan Bioquantum LS/967 energy filter (slit width of 20 eV) coupled to a Gatan K2 direct electron detector camera^23^. Automated data collection was performed with SerialEM using a beam-tilt data collection scheme^24^, acquiring one image per hole from 9 holes before moving the stage. Micrographs were recorded in super-resolution mode at a 165,000x magnification giving a pixel size of 0.4135 Å with defocus ranging from −0.8 to −3.5 µm. In total, 16,498 movies with 40 frames per movie were collected with a total exposure of 50 e^-^/Å^2^ (**Supplementary Table 2**).

### Image processing

Movie drift correction was performed in Motioncor2 using all frames, applying gain reference and cameras defect correction^25^. Image were binned twice, resulting in 0.826 Å/pixel size. Further initial image processing steps were performed in cryoSPARC v2.14.2^26^. CTF parameters were determined using “patch CTF estimation” on non-dose weighted micrographs. Realigned micrographs were then manually inspected using “Curate exposure” and low-quality micrographs were manually discarded for further image processing resulting in a curated 16,015 micrographs dataset (**Supplementary Fig. 2c**). LACV-L FL particles were then picked with “blob picker” using a circular particle diameter ranging from 90 to 150 Å, manually inspected and selected using “inspect particle picks”, extracted from dose weighted micrographs using a box size of 300*300 pixels^2^, resulting in a 4,065,475 particles dataset. Successive 2D classification run were used to eliminate bad quality particles displaying poor structural features resulting in 2,279,573 particles suitable for further image processing (**Supplementary Fig. 2b**). Per particle CTF was calculated. Subsequent steps were all performed in Relion 3.1^27, 28^. The entire dataset was divided in 4 (∼570,000 particles per subset) and subjected to 3D classification with coarse image alignment sampling (7.5°) using a circular mask of 170 Å and 10 classes (labelled “1^st^ 3D classification”, **Supplementary Fig. 2c**). LACV-L FL classes displaying a stable C-terminal conformation were kept and merged for further classification resulting in 566,025 selected particles (dotted squared, green maps in **Supplementary Fig. 2c**). Selected particles were subjected to a 2^nd^ run of 3D classification using finer angular sampling (3.7°) with a circular mask of 170 Å and was restricted to 10 classes (labelled “2^nd^ 3D classification”, **Supplementary Fig. 2c**). Particles from three 3D classes that display stable C-terminal regions without neighboring particles were selected to perform further high-resolution analysis, resulting in a 370,497 particle dataset. A previously obtained 3D class from the 2^nd^ run of 3D classification was low pass filtered to 15Å, extended by 10 pixels with a soft edge of 5 pixels and used for a final 3^rd^ run of 3D classification using finer angular sampling (1.8°) (labelled “3^rd^ 3D classification”, **Supplementary Fig. 2c**). Two classes (in dark blue, 59,152 particles) displayed a ten-base pair template-product RNA duplex in the LACV-L FL active site cavity and mimic an elongation state and one class (in cyan, 57,660 particles) displays a typical pre-initiation state with both 5′ 1-10 and 3′ 1-16 / 5′ 9-16 promoters bound. Both LACV-L FL pre-initiation and elongation-mimicking state subsets were submitted to 3D auto-refine using previous initial mask giving reconstructions at respectively 3.13 Å and 3.17 Å resolution using the FSC 0.143 cutoff criteria before post-processing. Masks to perform sharpening were generated using 3D refined maps low-pass filtered at 10 Å, extended by 4 pixels with 8 pixels of soft-edge. Post-processing was done using an applied B-factor of −40 Å^2^ and resulted in a map at 3.02 Å resolution for the elongation state and 3.06 Å resolution for the pre-initiation state using the FSC 0.143 cutoff criteria (**Supplementary Fig. 2c, Supplementary Fig. 3a**).

In order to deal with the high mobility and the small size of the CBD within LACV-L FL particles, the following advanced strategy was applied. 131,058 particles displaying a stable CBD (corresponding to its extreme position 1), originating from 4 out of the 10 classes from the 3^rd^ 3D classification (in orange in “3^rd^ 3D classification – CBD view”, **Supplementary fig. 2c**) were submitted to 3D auto-refine in order to get the best global accuracy alignment. A mask excluding LACV-L_46-1751_ residues, low pass filtered to 10Å, resampled and extended of 4 pixels with a soft edge of 6 pixels was then used for signal subtraction followed by particles re-centering on the mask center-of-mass. The resulting subtracted particles containing Endo_1-45_-Mid-CBD densities were subjected to 3D masked auto-refine with local angular searches. The obtained map was sharpened with an applied B-factor of −90 Å^2^ and resulted in a 3.54 Å resolution map according to the FSC 0.143 cutoff criteria.

A similar strategy was applied to deal with the ZBD flexibility. 287,363 particles displaying an ordered ZBD, originating from 7 out of the 10 classes from the 3^rd^ 3D classification (in red in “3^rd^ 3D classification – ZBD view”, **Supplementary fig. 2c**) were submitted to 3D auto-refine in order to get the best global accuracy alignment. A mask excluding LACV-L_1-1751/1841-1983_ residues, low-pass filtered to 10 Å, resampled and extended of 4 pixels with a soft edge of 6 pixels was used for signal subtraction followed by particles re-centering on the center-of-mass of this mask. Resulting subtracted particles containing Mid-ZBD densities were classified without alignment in order to detect potential heterogeneity. The most stable subset containing 131,396 particles was subjected to 3D masked auto refine with local angular searches. The resulting map was post processed with an applied B-factor of −90Å^2^, which result in a 3.4 Å resolution map according to the FSC 0.143 cutoff criteria.

For each final map, local resolution variations were estimated in Relion 3.1 (**Supplementary Fig. 3**). The 3D variability analysis was performed in cryoSPARC filtering resolution to 4 Å and using 3 modes.

### Model building in the cryo-EM maps

All the cryo-EM maps, namely pre-initiation map, elongation-mimicking map, CBD-mid domain map and ZBD-mid domain map were superimposed using Chimera^29^ previous to model building. The partial model determined in the 4.0 Å X-ray structure was used as a starting point to manually build into the cryo-EM maps using COOT^30^. The map chosen for model building was the one corresponding to the best resolution in the region built. As a result, the pre-initiation and elongation-mimicking maps were used to build the LACV-L core, the endonuclease domain, the mid domains, the CBD-mid domain map was used to build the CBD and the ZBD-mid domain map was used to build the ZBD. The sequence of the RNA duplex visible in the active site cavity at elongation state was deduced based on purines and pyrimidines, clearly visible in the density. As the 3′ and 5′ vRNA incubated respectively contain 16 and 10 nucleotides, 6 nucleotides of the vRNA should be present in the product exit tunnel and none in the template exit tunnel. Blurred density is visualized in the product exit tunnel due to the large flexibility of the nucleotides. Blurred RNA density is also visible in the template exit tunnel suggesting that some particles have encapsidated the RNA in different positions than the one shown in **Fig. 1c, 4d**. After initial manual building in COOT, the models were iteratively improved by Phenix-real space refinement^31^ and manual building in COOT. Validation was performed using the Phenix validation tool, and model resolution was estimated at the 0.5 Fourier Shell Correlation (FSC) cutoff. Figures were generated using ChimeraX^32^

### Multiple alignment

6 sequences of major Peribunyaviruses (L_BUNYW for Bunyamwera virus, L_SBVBG for Schmallenberg virus, L_MCAV for Macaua virus, L_WBV for Wolkberg virus, L-Oya for Oya virus) were aligned using MUSCLE^33^ and presented with ESPript^34^.

**Table 2.** Crystallographic statistics

| Data Collection | LACV L FL<br>5' hook<br>3' template<br>5' duplex region |
| --- | --- |
| Beamline (ESRF) | ID29 |
| Wavelength (Å) | 0.9724 |
| Space group | C2 |
| Unit cell: a, b, c (Å) | a=371.19, b=145.32,<br>c=234.19, β=116.33 |
| Resolution (Å) | 172.95-3.96<br>(4.35-3.96) |
| R<sub>meas</sub> | 0.163 (1.285) |
| I/σI | 6.5 (1.6) |
| CC(1/2) | 0.995 (0.47) |
| Completeness (%) |  |
| Spherical | 60.8 (12.4) |
| Ellipsoidal | 91.3 (55.8) |
| Redundancy | 11.6 (13.3) |
| **Refinement** |  |
| No. reflections work/free | 57697/1514 |
| R<sub>work</sub> | 0.265 (0.328) |
| R<sub>free</sub> | 0.302 (0.323) |
| No. protein atoms | 34547 |
| Average B-factor (Å<sup>2</sup>) | 188.9 |
| R.M.S. deviations |  |
| Bond lengths (Å) | 0.002 |
| Bond angles (°) | 1.153 |
| **Validation** |  |
| Ramachandran favoured | 91.1 |
| Ramachandran outliers | 1.32 |
| Molprobity score | 2.33 |
| Clash score | 5.24 |

## Data availability

Coordinates and structure factor have been deposited in the Protein Data Bank and the Electron Microscopy Data Bank.

LACV-L pre-initiation complex (X-ray crystallography) PDB XXXX

LACV-L pre-initiation complex PDB XXXX EMDB XXXX

LACV-L elongation complex PDB XXXX EMDB XXXX

LACV-L CBD and mid domain map EMDB XXXX

LACV-L ZBD and mid domain map EMDB XXXX

## ACKNOWLEDGEMENTS

We thank Karine Huard, Angélique Fraudeau and Alice Aubert for technical advices on expression and purification; Ambroise Desfosses, Leandro Estrozi and Alister Burt for discussion on image processing; Aymeric Peuch for help with the usage of the EM computing cluster; Dominique Housset for general discussion on the project and Irina Gutsche for support.

This work used the platforms of the Grenoble Instruct-ERIC center (ISBG ; UMS 3518 CNRS-CEA-UGA-EMBL) within the Grenoble Partnership for Structural Biology (PSB), supported by FRISBI (ANR-10-INBS-05-02) and GRAL, financed within the University Grenoble Alpes graduate school (Ecoles Universitaires de Recherche) CBH-EUR-GS (ANR-17-EURE-0003).The electron microscope facility is supported by the Auvergne-Rhône-Alpes Region, the Fondation Recherche Médicale (FRM), the fonds FEDER and the GIS-Infrastructures en Biologie Santé et Agronomie (IBISA). We acknowledge the European Synchrotron Radiation Facility (ESRF) for provision of beam time on CM01. We thank all platform staff that enabled us to perform these analyses. This work was supported by the IDEX IRS G7H-IRS17H50 and the ANR-19-CE11-0024-02 to HM.

## AUTHOR CONTRIBUTIONS

B.A., P.G. and J.R. expressed and purified LACV-L FL. P.G. crystallized LACV-L FL. S.C. P.G. and J.R. solved the X-ray structure. S.C. built an initial model based on X-ray crystallography data. B.A. prepared cryo-EM grids. B.A. and H.M. collected cryo-EM data on a Thermofischer Glacios EM thanks to advices and training from G.S. who set up and maintains the IBS EM platform. G.E. set up SerialEM data collection scheme and collected the high-resolution cryo-EM dataset on the Thermofischer Krios EM at ESRF (CM01). B.A. performed advanced cryo-EM image processing and 3D reconstructions. H.M., B.A. and S.C. built the models based on the cryo-EM maps and performed structural analysis. H.M. and G.S. co-supervise B.A ; J.R. and S.C. co-supervised P.G. The project was conceived by H.M. and S.C. This project used funding obtained by H.M., G.S. and S.C. The manuscript was written by H.M. and B.A. with input from all authors.

---

**Table supplement 2:** Cryo-EM data collection, refinement and validation statistics

|  |  |  |  |  |
| --- | --- | --- | --- | --- |
| **Data Collection** |  |  |  |  |
| Microscope | ThermofischerTitan Krios |  |  |  |
| Camera | Gatan K2 Summit + GIF |  |  |  |
| Magnification | 165000 |  |  |  |
| Voltage (kV) | 300 |  |  |  |
| Number of frames | 40 |  |  |  |
| Electron exposure total (e<sup>-</sup> /Å<sup>2</sup>) | 50 |  |  |  |
| Defocus range (μm) | 0.8-3.5 |  |  |  |
| Pixel size (Å) | 0.4135 in super-resolution, 0.827 used for processing |  |  |  |
| Symmetry imposed | C1 |  |  |  |
| Initial/Final micrographs (no.) | 16498/16015 |  |  |  |
| **Processing** | **Pre-**<br>**initiation**<br>**(ENDO,**<br>**CORE, MID,**<br>**ZBD β-**<br>**hairpin**<br>**built, RNA)** | **Elongation**<br>**(ENDO,**<br>**CORE, MID,**<br>**ZBD β-**<br>**hairpin**<br>**built, RNA)** | **CBD** | **ZBD without**<br>**β-hairpin** |
| Final particles (no.) | 57660 | 59152 | 131058 | 287363 |
| Map resolution (Å) 0.143 FSC<br>threshold | 3.06 | 3.02 | 3.54 | 3.39 |
| Map resolution range (Å) | 2.9-5 | 2.9-5 | 3.5-4.3 | 3.3-4.4 |
| **Refinement** |  |  |  |  |
| Initial model used | 5AMQ | 5AMQ | - | - |
| Model resolution (Å) 0.5 FSC threshold | 3.06 | 3.12 | 4.03 | 3.53 |
| Map sharpening B factor (Å<sup>2</sup>) | -40 | -40 | -90 | -90 |
| Model composition |  |  |  |  |
| Protein residues | 1815 | 1831 | 129 | 198 |
| Ligands | 29 | 46 | 0 | 1 |
| B-factor (Å<sup>2</sup>) |  |  |  |  |
| Protein<br>min-max (mean) | 12.93-145.07<br>(55.74) | 40.59-158.48<br>(74.67) | 69.72-127.37<br>(93.35) | 51.01-129.50(77.7<br>9) |
| R.m.s deviations |  |  |  |  |
| Bond lengths (Å) | 0.008 | 0.009 | 0.005 | 0.006 |
| Bond angles (°) | 0.889 | 0.931 | 0.924 | 0.828 |
| **Validation** |  |  |  |  |
| MolProbity score | 1.46 | 1.54 | 1.88 | 1.62 |
| Clashscore | 3.57 | 3.79 | 4.97 | 3.99 |
| Poor rotamers (%) | 0.24 | 0.53 | 0.00 | 0.00 |
| Ramachandran plot |  |  |  |  |
| Favored (%) | 95.53 | 94.57 | 87.07 | 93.40 |
| Allowed (%) | 4.47 | 5.43 | 12.97 | 6.60 |
| Disallowed (%) | 0 | 0 | 0 | 0 |

**Supplementary figure 1:**
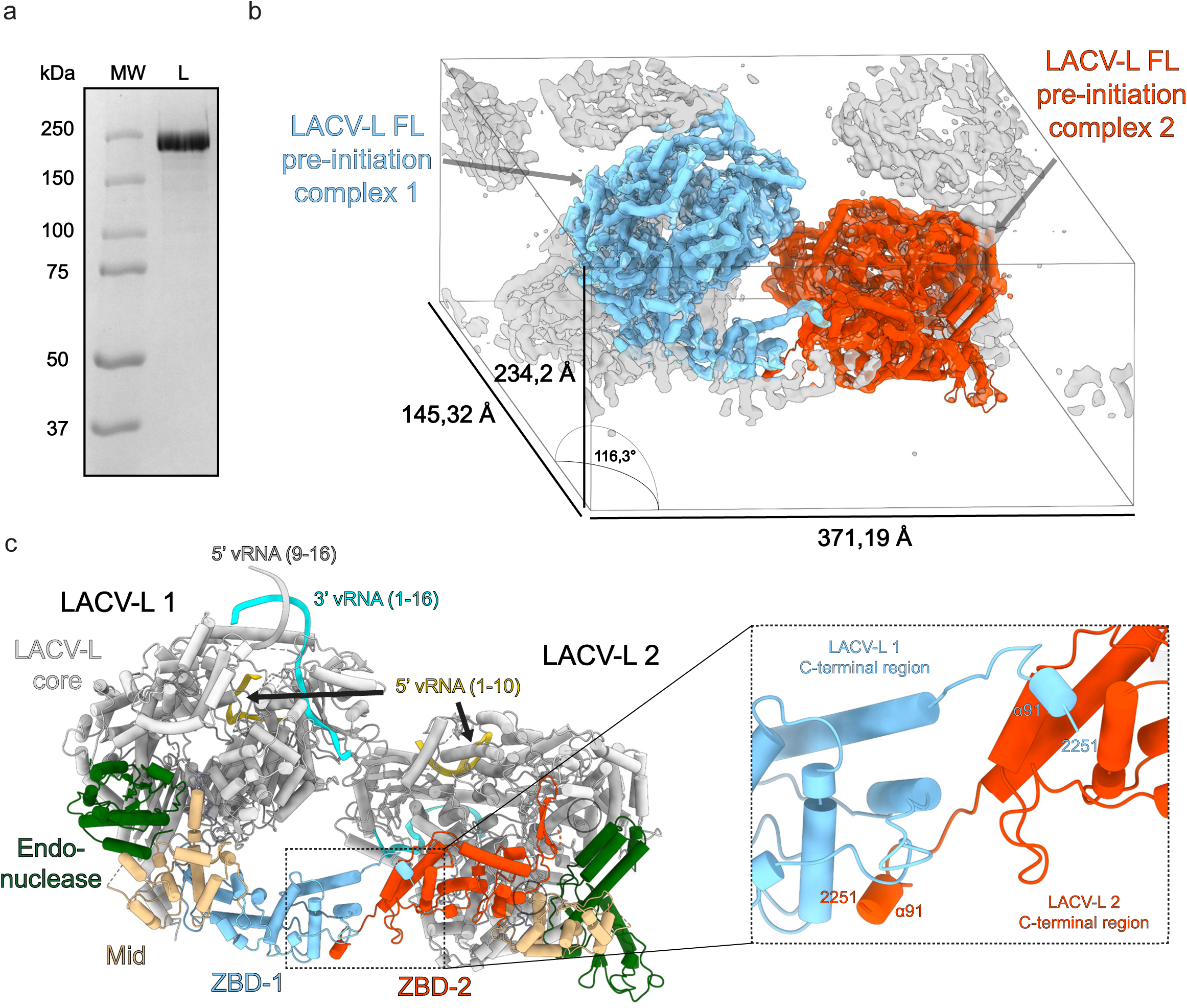
Purification and X-ray crystal structure. **a**, 8% SDS-PAGE gel of LACV-L FL purification after gel filtration step (MW: Molecular weight; L: LACV-L FL). **b**, Electron density map of one asymmetric unit. The two polymerase complexes at pre-initiation are respectively colored in blue, red and labelled. Asymmetric unit dimension and angles are displayed. **c**, X-ray crystallography structure shown in the same orientation as in b displaying the 2 polymerase complexes of the asymmetric unit. The core regions are colored in white, the endonucleases in green, the mid domains in beige and the ZBD according to LACV-L 1/LACV-L 2 colors in blue and orange. The swap of the last α-helix 91 is shown in a close up view and the C-terminal residue 2251 is labelled. The RNA promoter positions are shown.

**Supplementary figure 2:**
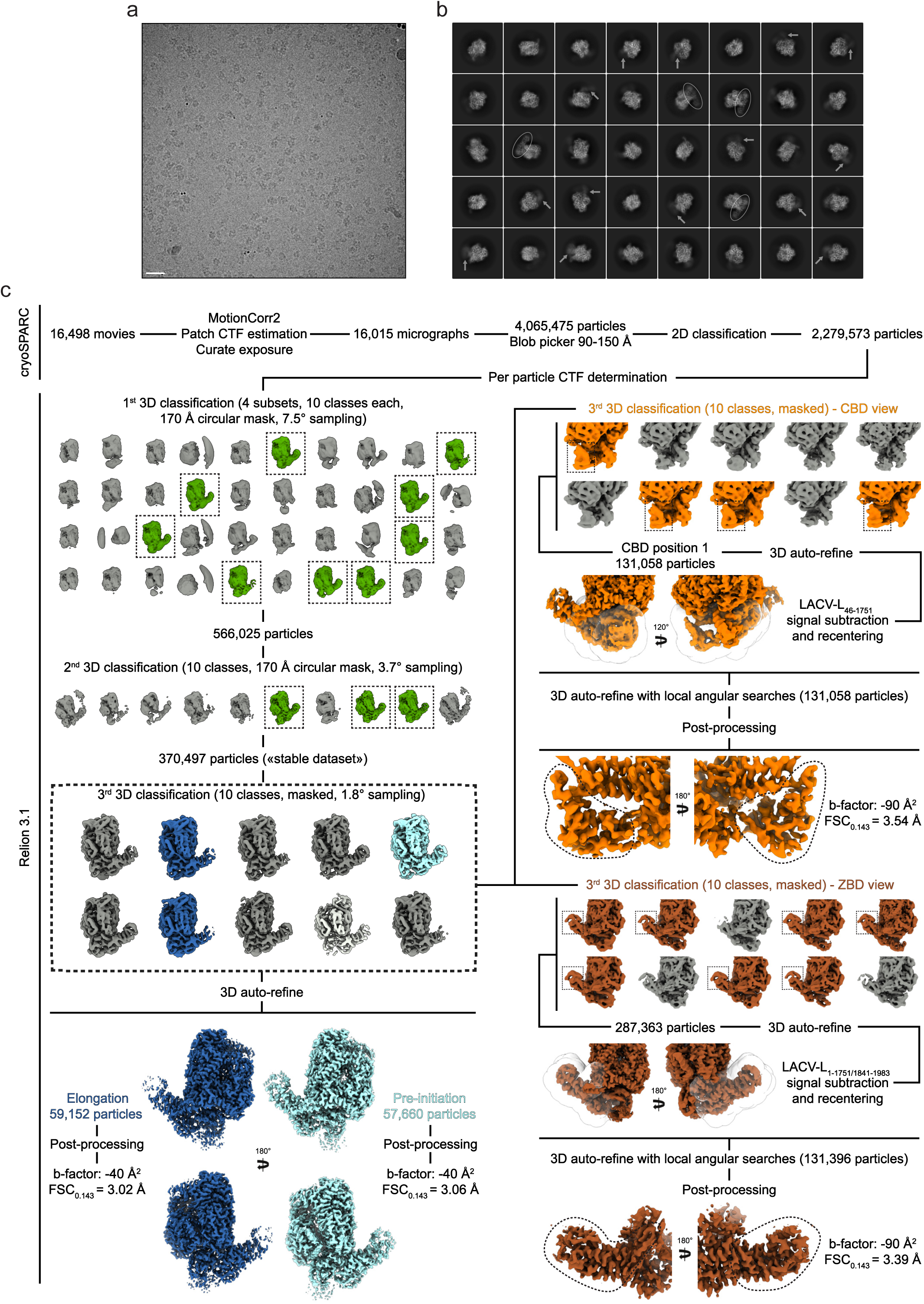
Cryo-EM data collection and image processing. **a**, Cryo-electron micrograph of LACV-L FL collected on the ESRF CM01 Titan Krios equipped with a K2 direct electrons detector camera at −2 μm defocus. Scale bar = 200 Å. **b**, Representative 2D classes of LACV-L FL. Flexibility of the LACV-L C-terminal region seen in some class averages is highlighted with white arrows. Stabilized LACV-L C-terminal regions visible in other class averages are surrounded by a white circle. **c**, Cryo-EM image processing pipeline is described in detail in the “Image processing” method section.

**Supplementary figure 3:**
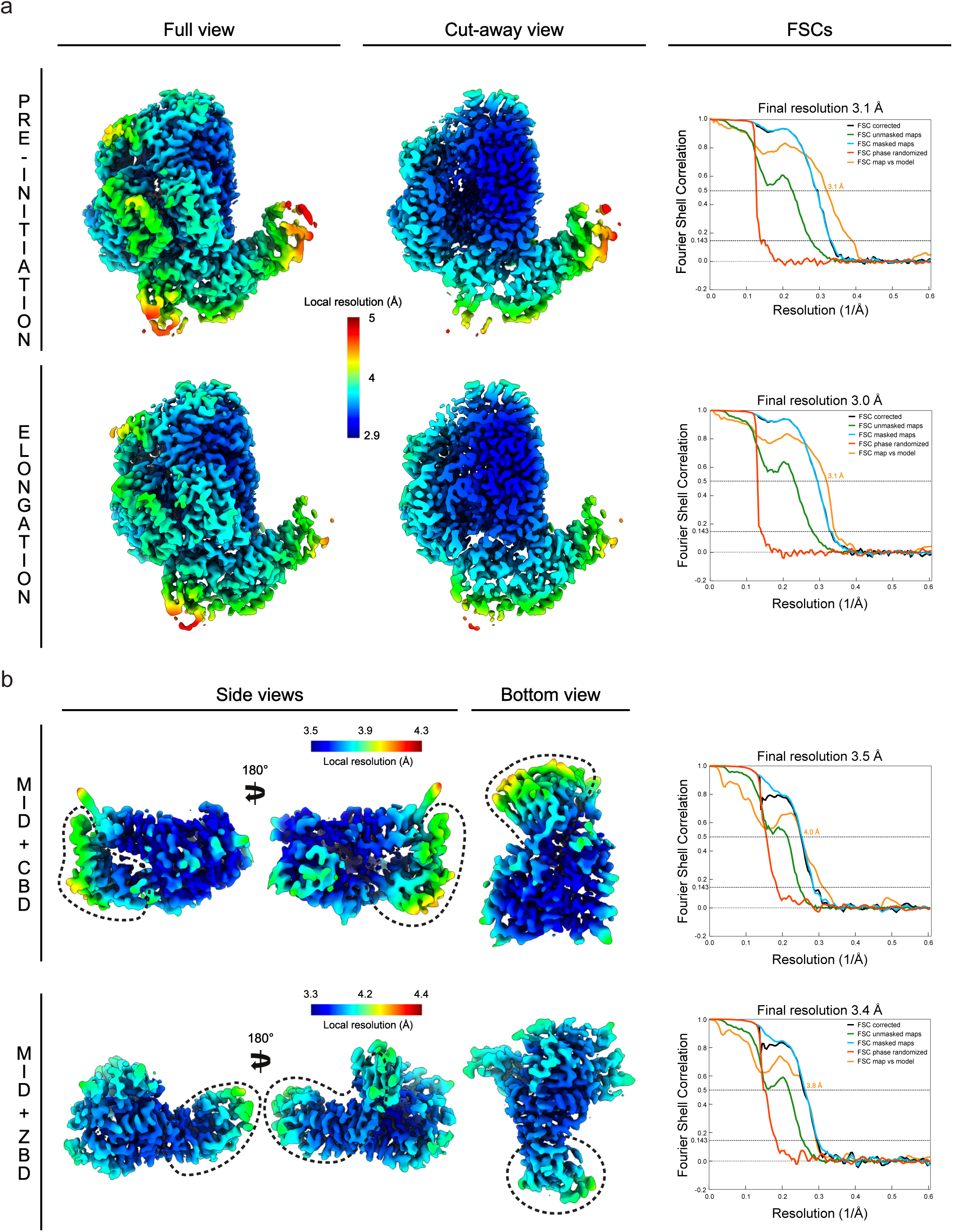
Cryo-EM maps local resolution and FSCs. **a**, Full view and cut-away view of the pre-initiation and elongation-mimicking cryo-EM maps. Maps are filtered and colored according to their local resolution. Resolution range coloring is indicated. FSCs corrected, FSCs of unmasked maps, FSCs of masked map and phase randomized FSCs are respectively displayed in dark blue, green, light blue and red. The FSC of the map and the model are displayed in orange. The model refined in the pre-initiation and the elongation-mimicking cryo-EM maps corresponds to the core, the mid domain, the endonuclease domain and the ZBD β-hairpin. The gold-standard Fourier shell correlation (FSC) of masked maps indicates a respective resolution of 3.1 Å and 3.0 Å for the pre-initiation and the elongation-mimicking cryo-EM maps with the FSC = 0.143 criteria. **b**, Side view and bottom view of the subtracted maps containing (i) the CBD and the mid domain, (ii) the ZBD and the mid domain. The maps are filtered and colored according to their local resolution. Resolution range coloring is indicated. The CBD position is surrounded by a dotted line in the map containing the CBD and the mid domain. The ZBD position is surrounded by a dotted line in the map containing the ZBD and the mid domain. FSCs are displayed and colored as in a. The gold-standard Fourier shell correlation of masked map indicates a resolution of 3.5 Å for the CBD-mid domain map and 3.4 Å resolution for the ZBD-mid domain map (FSC = 0.143 criteria).

**Supplementary figure 4:**
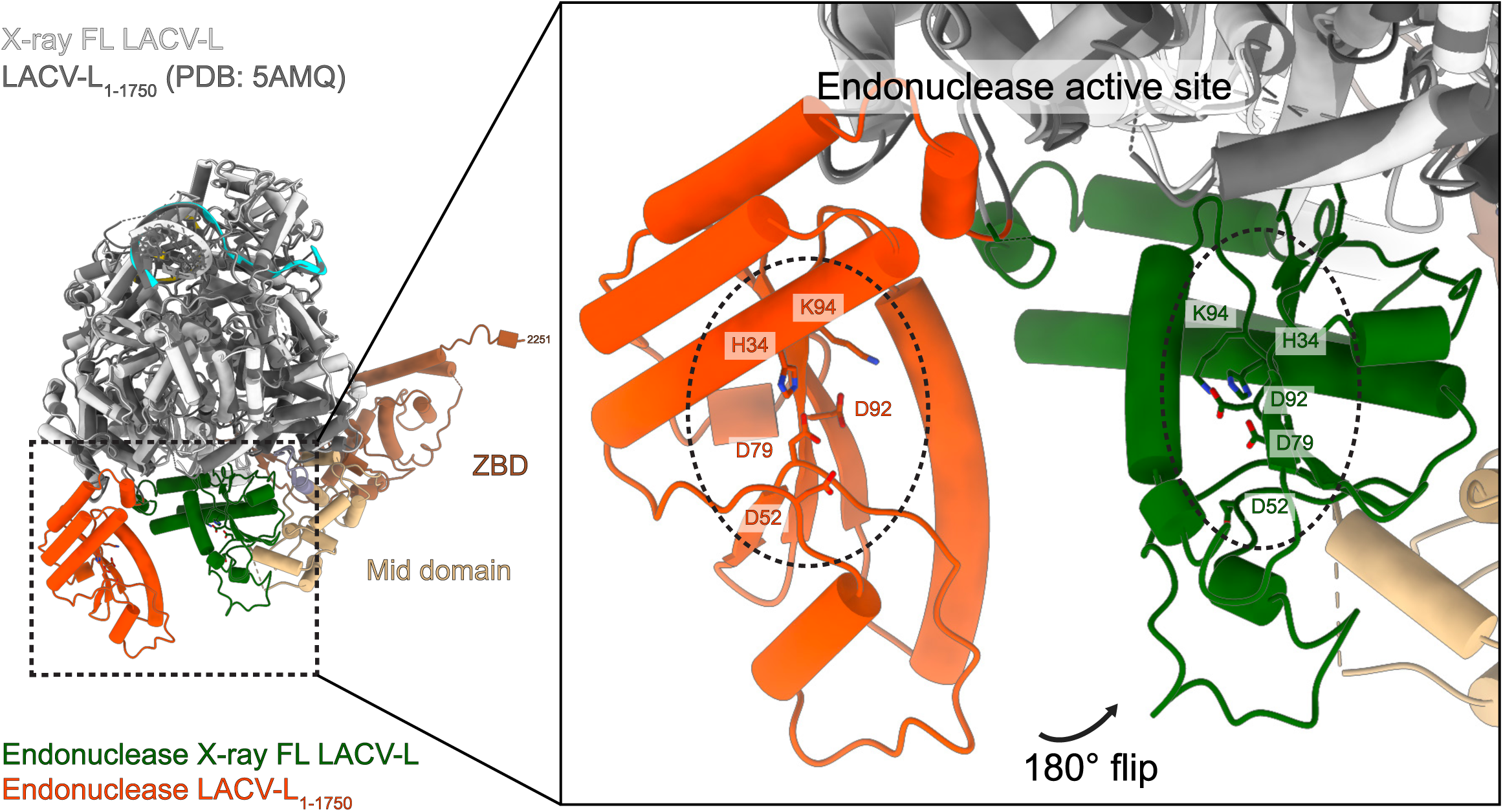
Endonuclease movement between LACV-L FL and LACV-L_1-1750_. Superimposition of LACV-L FL and LACV-L_1-1750_ (PDB: 5AMQ). LACV-L FL core is shown in light grey, LACV-L_1-1750_ core in light grey. LACV-L FL endonuclease is shown in green, LACV-L_1-1750_ endonuclease in red. The 180° rotation of the endonuclease domain is indicated. The endonuclease active site residues are displayed and are surrounded by a dotted ellipse on the close-up view.

**Supplementary figure 5:**
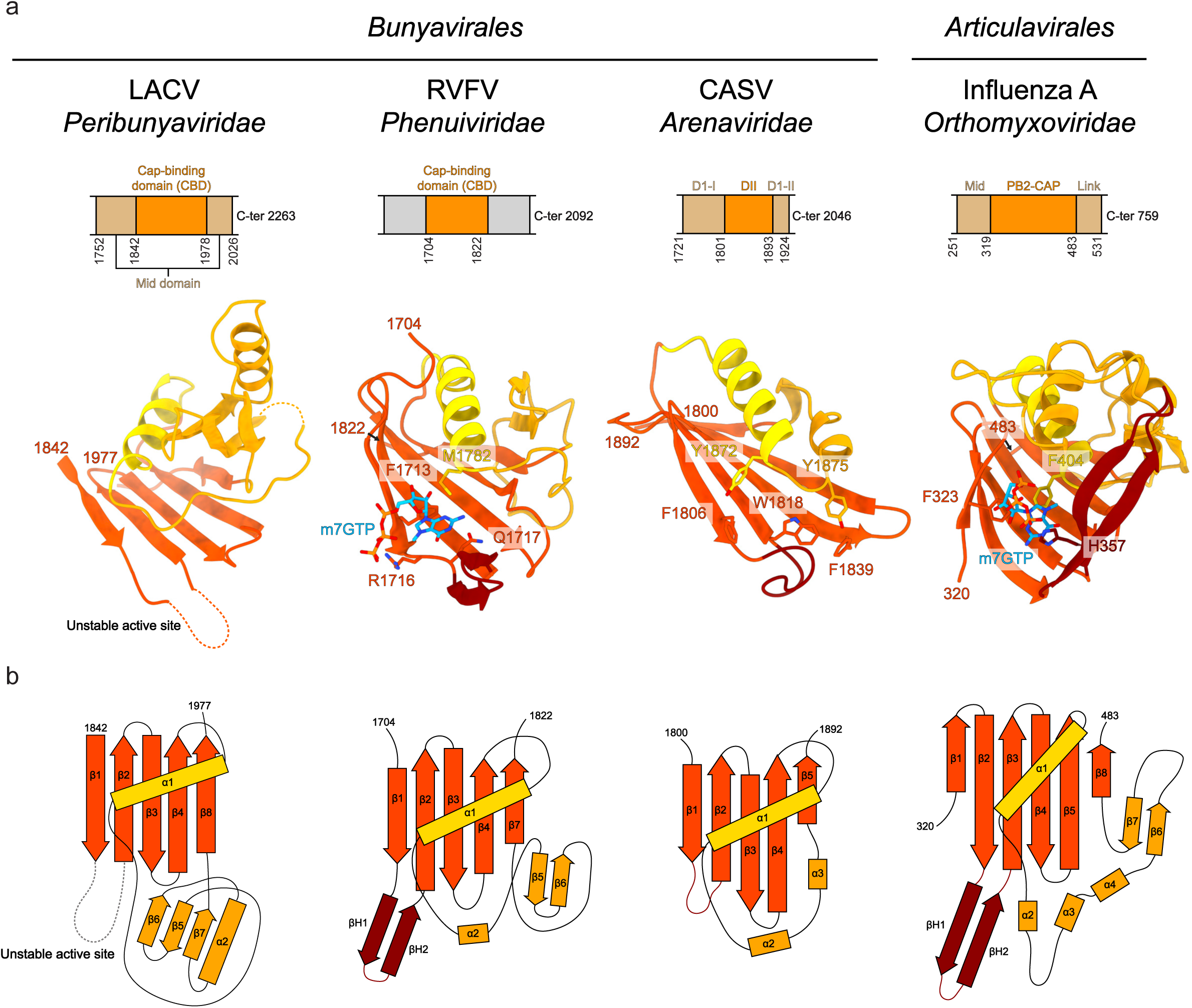
Structural comparison of sNSV CBD. **a**, Schematic representation of domains surrounding the CBD of LACV, RVFV, CASV and influenza A virus. Structures of LACV-L CBD, RVFV-L CBD (PDB: 6QHG), CASV-L putative CBD (PDB: 5MUZ) and influenza A PB2-CBD (PDB: 2VQZ) shown as ribbon. Similar structure elements are depicted in the same color: structurally conserved β-sheet in dark orange, α-helix in gold and specific insertions in orange. Residues involved in cap binding are shown and labelled. m^7^GTP molecules are colored in cyan. **b**, Schematic representation of CBDs shown as in **a**. LACV-L CBD loop suggested to contain the m^7^GTP binding site is shown as a dotted line. For clarity, secondary structures are numbered in the same way starting from α1 and β1.

**Supplementary figure 6:**
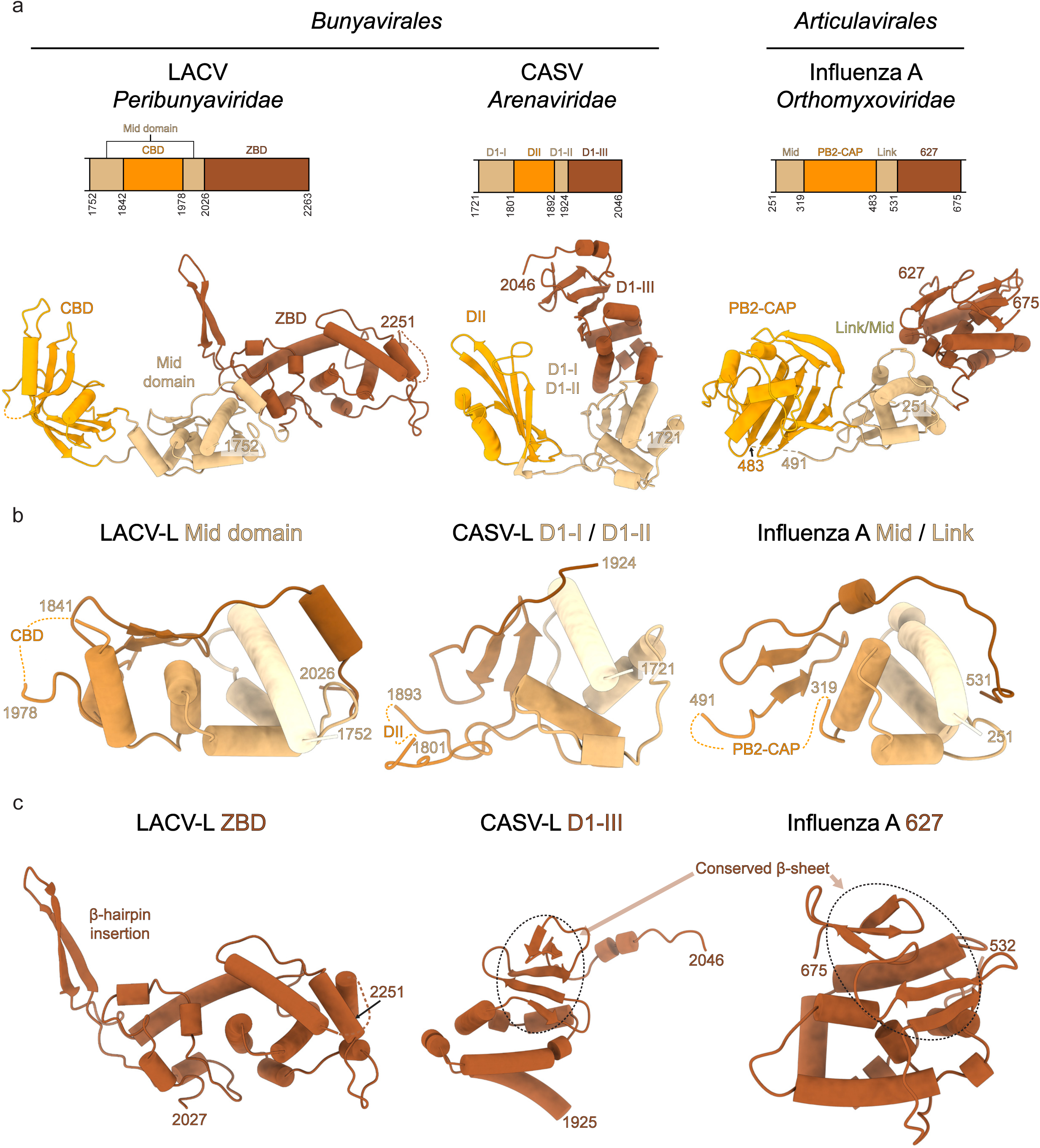
Comparison of C-terminal domain organization between sNSV polymerases. **a**, C-terminal domains of LACV-L, CASV-L (PDB: 5MUS) and influenza A virus polymerase (PDB: 4WSB): schematic representation (top) and structure (bottom). Similar domains are colored in the same way. **b**, Superimposed structures of LACV-L mid domain, CASV D1-I/D1-II and influenza A mid/link. Rainbow colors from beige to brown is used to color from the N-terminal to the C-terminal. **c**, Structure of LACV-L ZBD, CASV D1-III and influenza A 627. CASV D1-III and influenza A 627 domains are superimposed. Their conserved β-sheet is surrounded by a dotted line.

**Supplementary figure 7:**
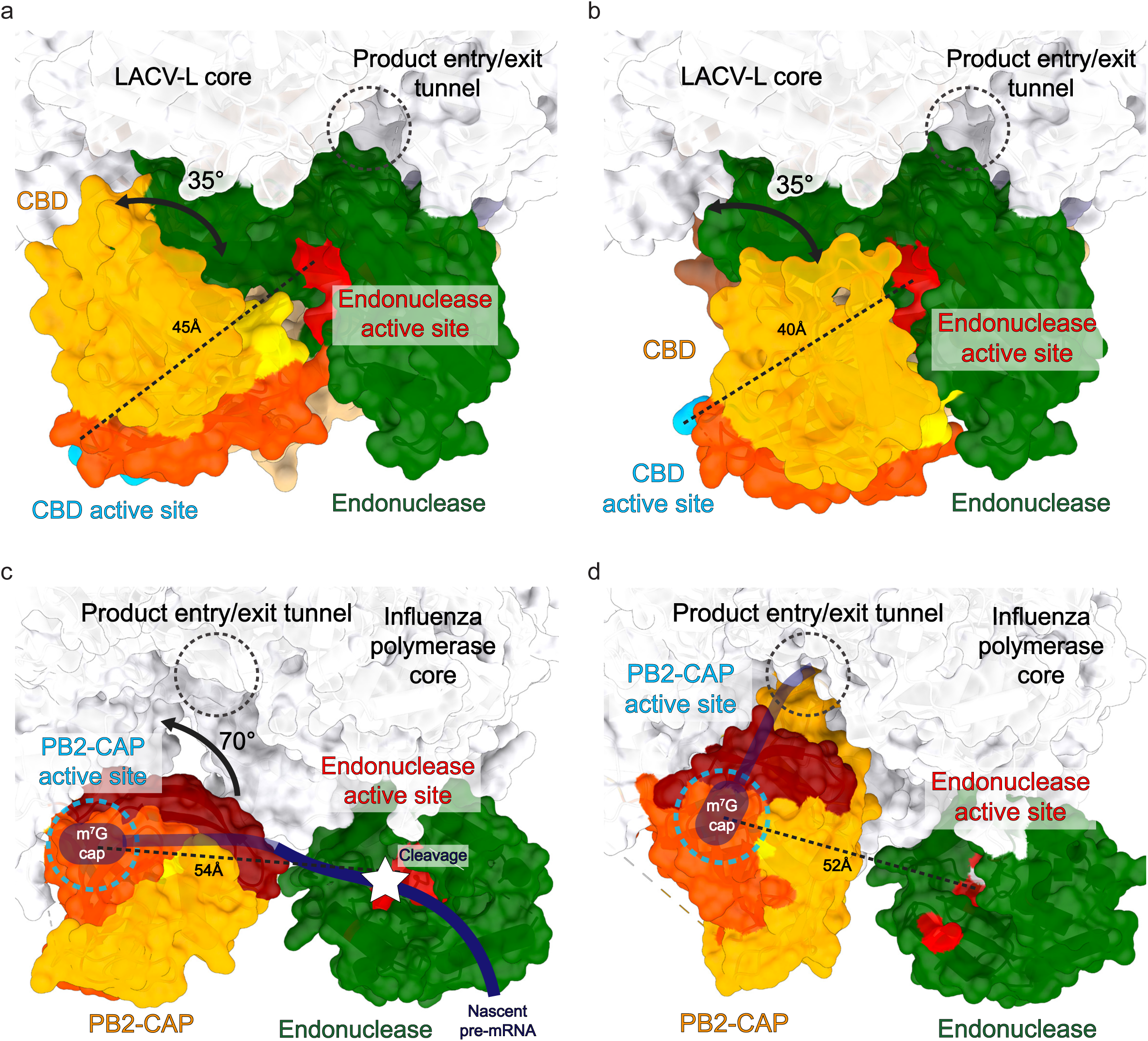
Comparison of CBD positions between LACV-L and influenza virus polymerase. **a**, LACV-L with CBD in extreme position 1. LACV-L core is shown in white, the endonuclease domain in green with the endo-nuclease active site in red. LACV-L CBD is colored as in Fig. 3 with the active site colored in cyan. The distance between the endo-nuclease and the CBD active sites is labelled. The product entry/exit tunnel position is shown with a dotted circle. The 35° movement of the CBD between extreme position 1 and 2 is shown. **b**, LACV-L with CBD in extreme position 2. Same colors and labels as in **a**. **c**, Influenza virus polymerase (PDB: 4WSB) with the cap-binding domain in a position compatible with binding and cleavage of the capped RNA primer (shown as a blue thick line, transparent when the binding site is not visible in the orientation chosen). RNA cleavage is indicated with a white star. The 70° rotation between PB2-CAP in **c** and **d** is shown. **d**, Influenza virus polymerase (PDB: 6QCT) with the cap-binding domain in a position enabling entry of the capped RNA primer in the product entry/exit channel. Color and labelling as in **c**.

**Supplementary figure 8:**
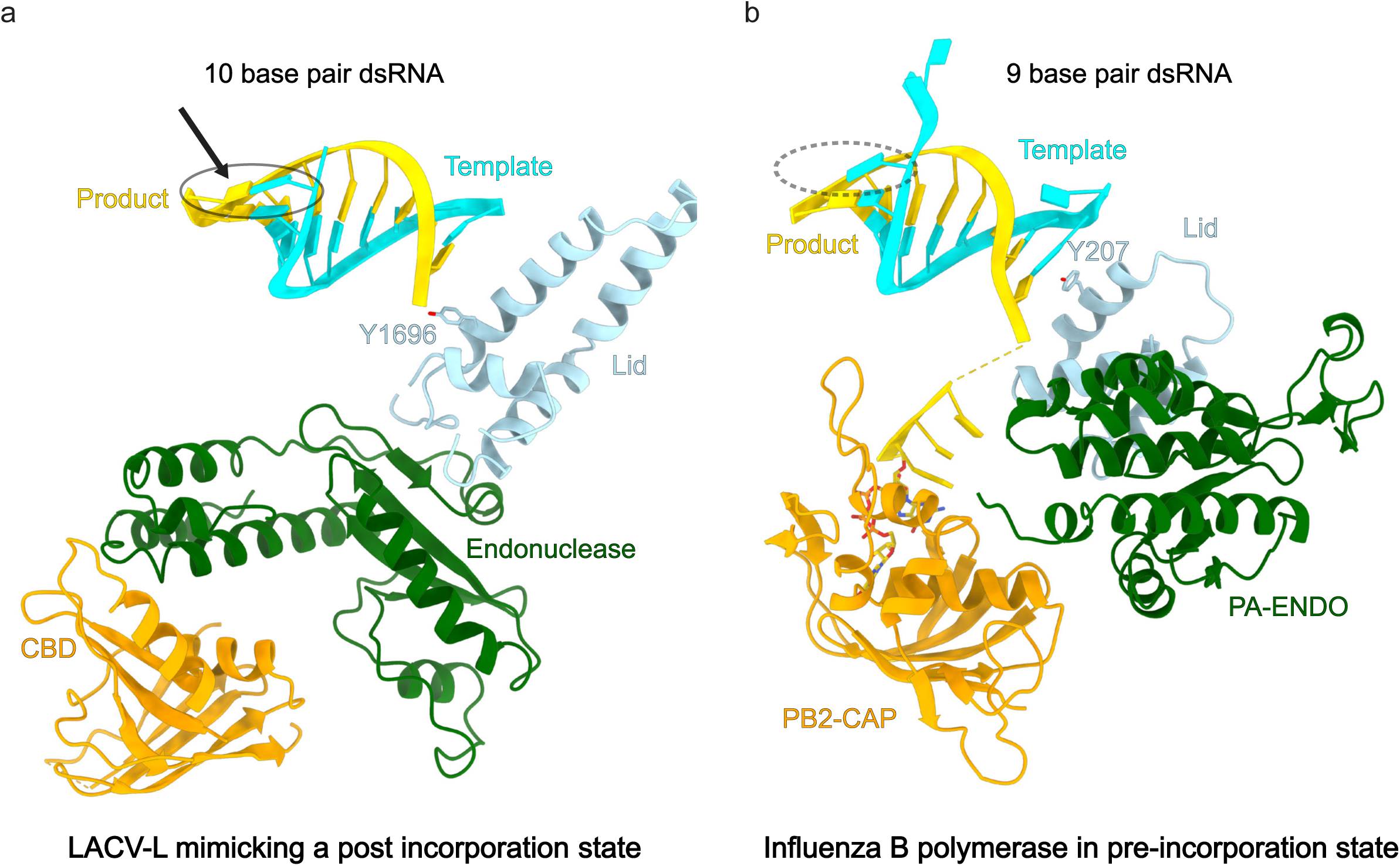
Comparison between LACV and influenza polymerase elongation states. **a**, **b**, The template, product, lid domain, endonuclease domain and CBD are respectively displayed in blue, yellow, light blue, green and orange. The nucleotide that mimics the last incorporated residue in LACV-L is shown with an arrow. The 10^th^ base-pair is surrounded with an ellipse. Equivalent missing base-pair in influenza polymerase (PDB: 6QCT) is shown with a dotted circle. Y1696 of LACV-L and Y207 of influenza PB2 that are lid residues preventing double-strand continuation are shown. LACV-L Y1696 interacts with the product RNA whereas influenza PB2 Y207 interacts with the template RNA.

**Supplementary figure 9:**
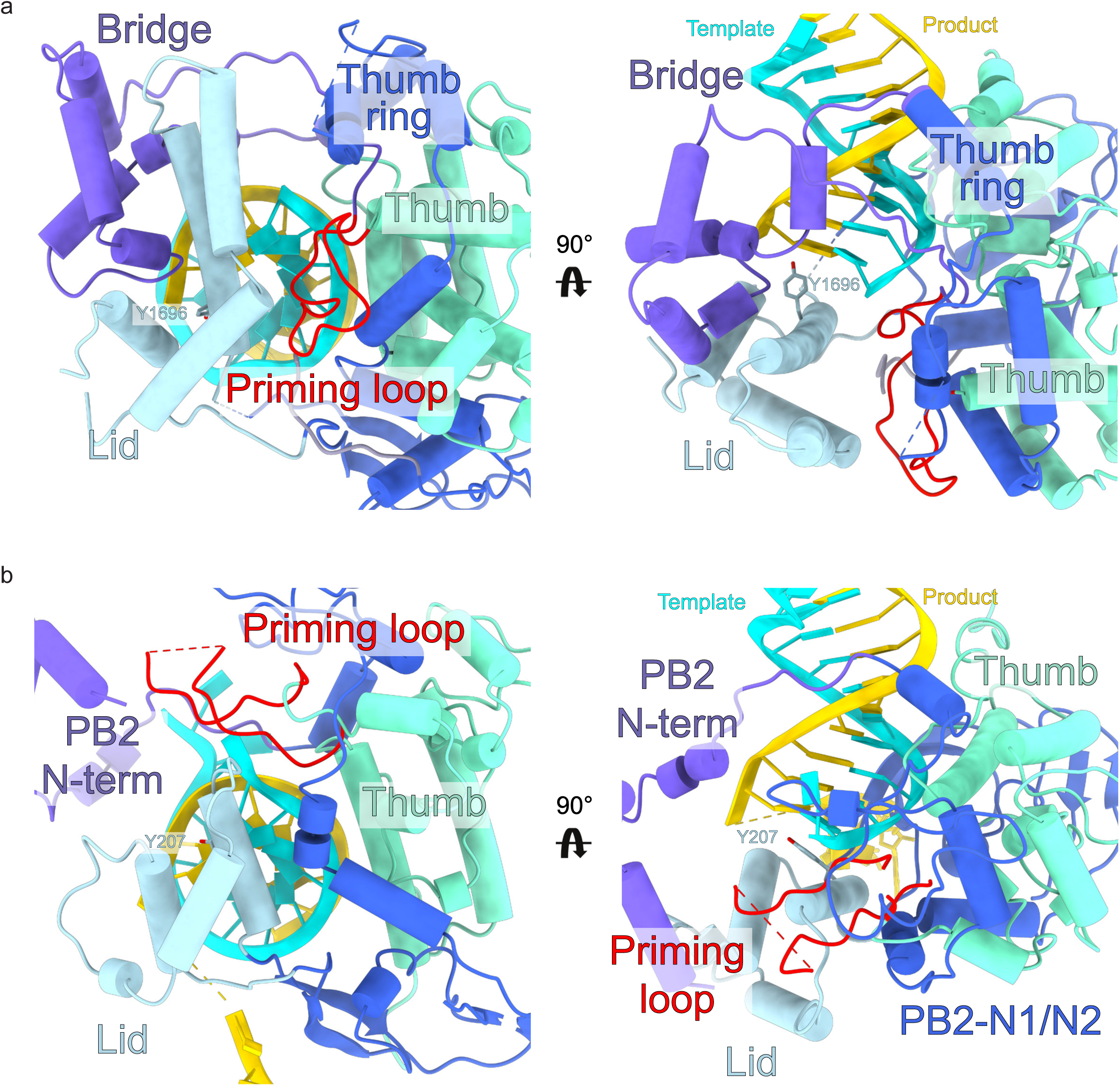
Comparison between LACV and influenza polymerase priming loop at elongation. **a**, LACV-L thumb, thumb ring, bridge and lid domains are displayed in turquoise, blue, purple and light blue respectively. The priming loop is shown in red. The template RNA is shown in cyan and the product in gold. dsRNA top view (left) and side view (right) are shown. **b**, Equivalent elements are shown using the same color code in influenza virus polymerase (PDB: 6QCT). The priming loop extremity is disordered and is shown as a dotted line.

**Supplementary Figure 10:**
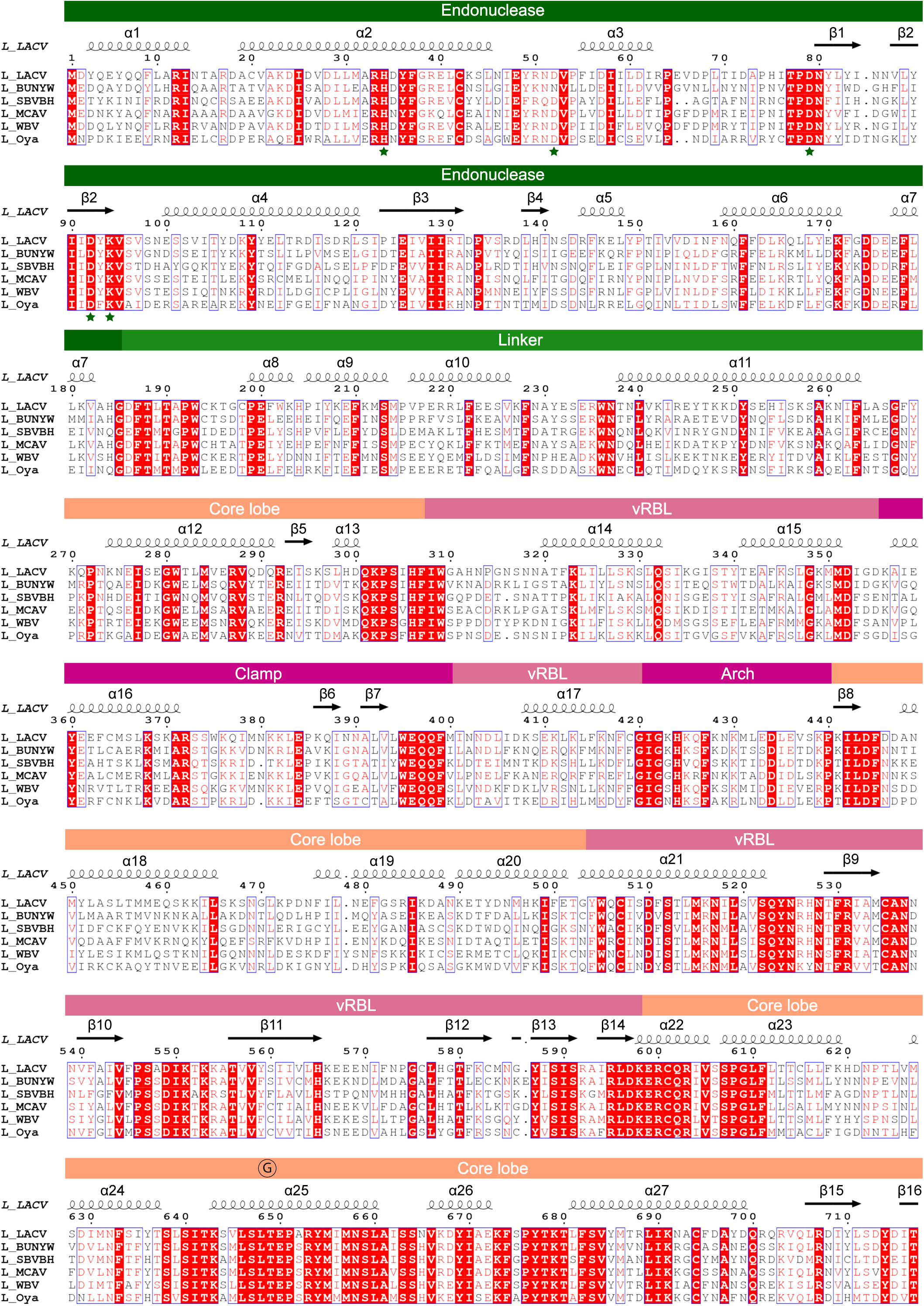
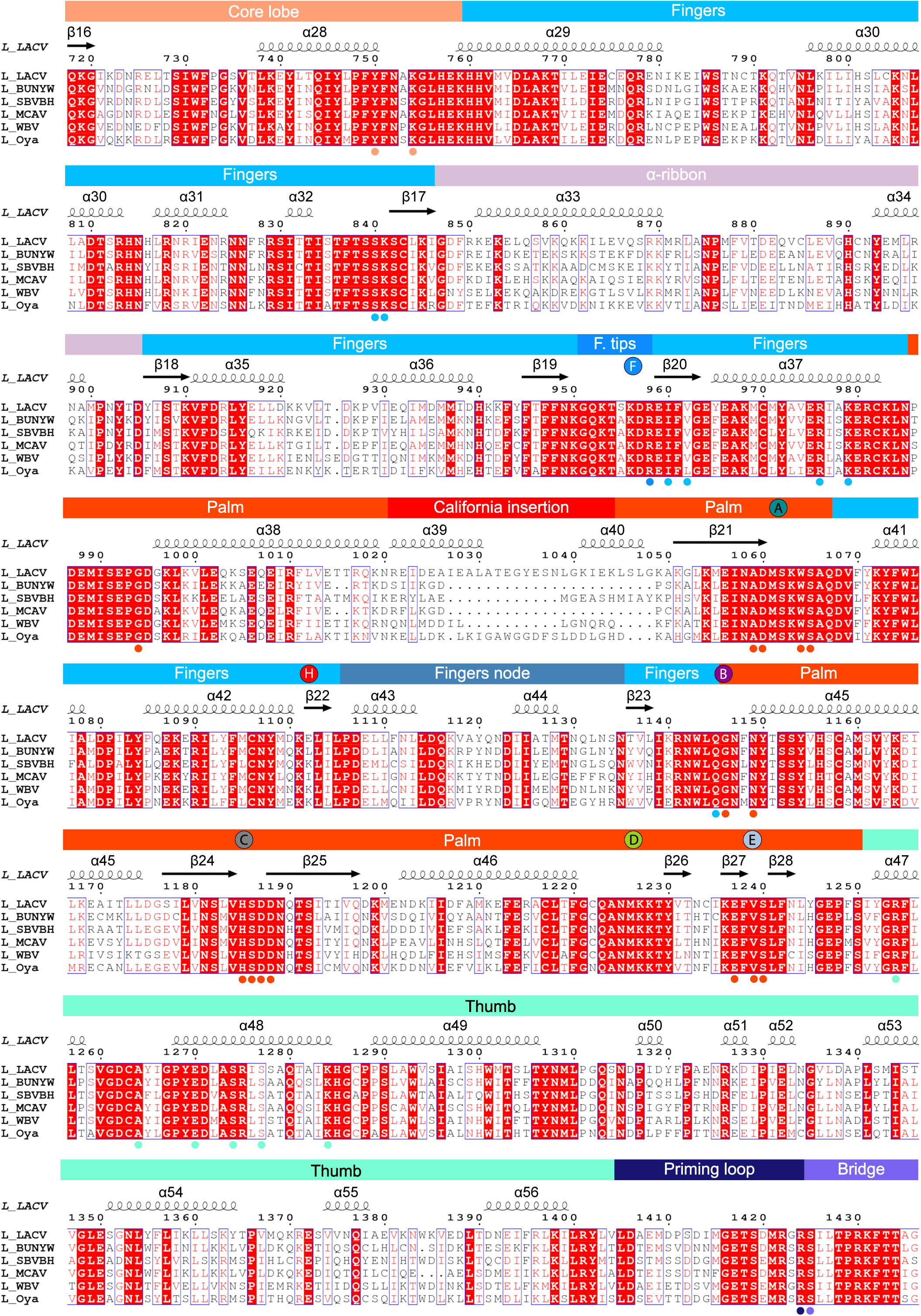
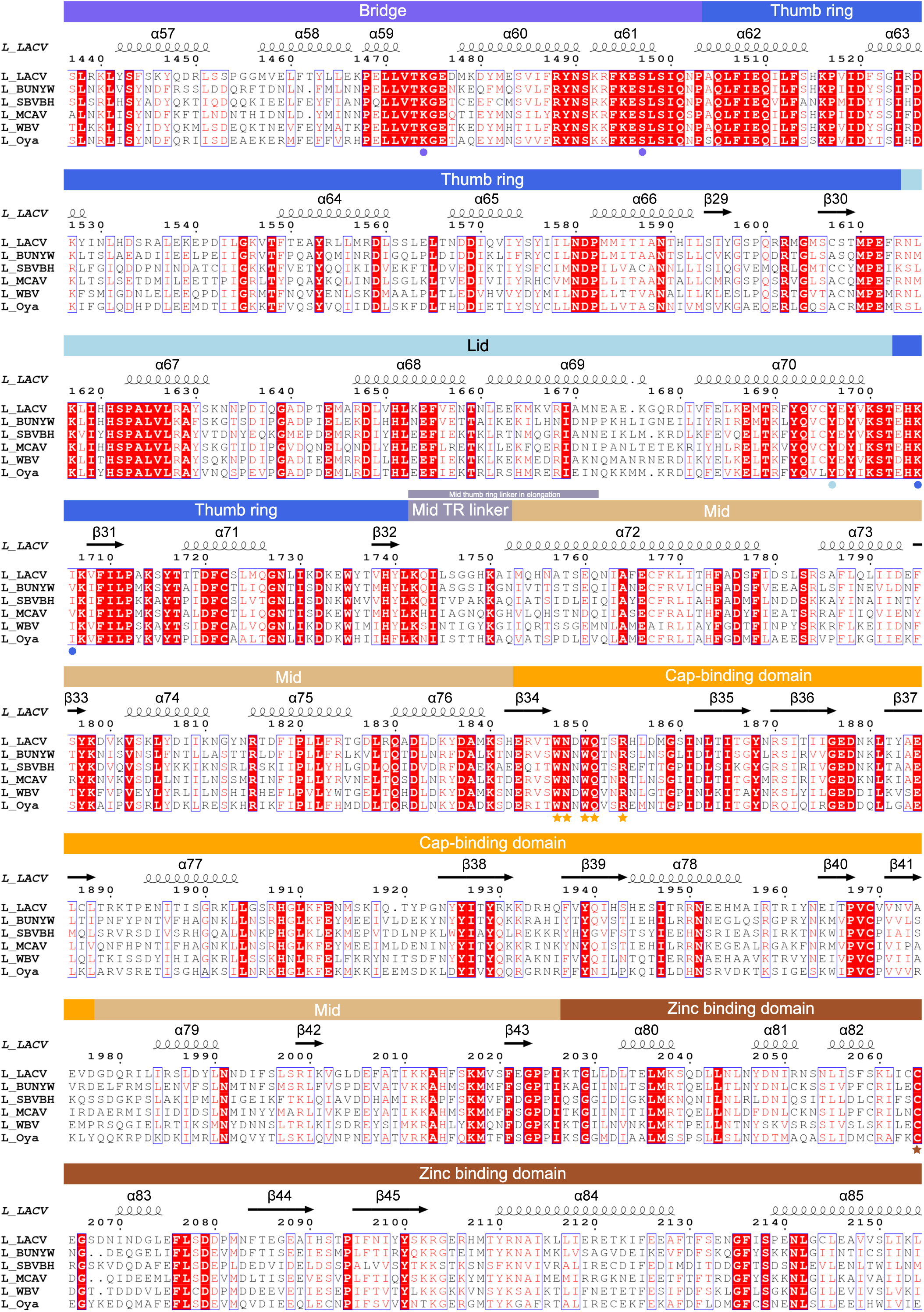
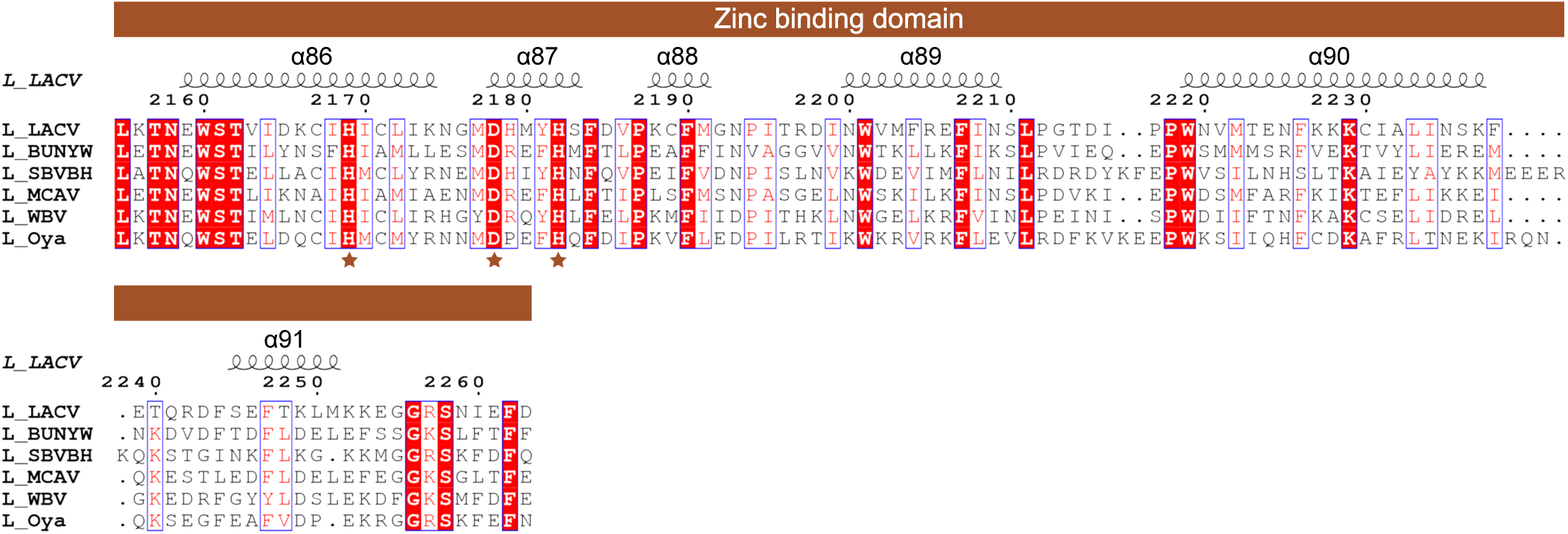
Multiple alignment of *Peribunyaviridae* L proteins. Multiple alignment of six *Peribunyaviridae* L proteins: LACV, Bunyamwera virus (BUNYW), Schmallenberg (SBVBG), Macaua virus (MCAV), Wolkberg virus (WBV) and Oya virus. LACV-L secondary structures are shown and numbered. Domain positions and motifs are indicated. Endonuclease and CBD active site residues are labelled with green and gold stars respectively. Residues that coordinate the zinc are shown with a brown star. Residues that interact with the template/product RNA are labelled with a circle colored based on their domain localization.

